# Ecoclimatic variables and karyotype features are poor predictors of neo-sex chromosome evolution in mammals

**DOI:** 10.1101/2025.11.05.686331

**Authors:** Jimmy Choi, Polly Campbell, Jonathan J. Hughes

## Abstract

The XX/XY chromosome system is highly conserved in therian mammals. Despite this evolutionary persistence, multiple variant sex chromosome systems exist in mammals. The potential contribution of genomic conflict to the evolution of neo-sex chromosomes has been studied extensively, while the influence of climate and ecology on the emergence and persistence of neo-sex chromosomes is less well understood. Building on previous literature that suggests that ecoclimatic factors can influence intralocus sexual conflict and other forms of genome evolution, we tested whether neo-sex chromosomes in mammals are associated with these factors. We conducted a phylogenetic logistic regression in three representative mammal families using a dataset of variant mammalian sex chromosomes and a database of tetrapod traits and environmental associations. We also incorporated a dataset of mammal karyotypes to test for potential signatures of meiotic drive. When using a single phylogeny, none of the seven ecoclimatic variables tested nor chromosome morphology were significantly correlated with neo-sex chromosomes. However, correlations between neo-sex chromosomes and both range size and mean annual temperature were data-dependent and sensitive to differences in tree topology, demonstrating the importance of accounting for alternative evolutionary hypotheses in comparative methods.

## Introduction

The XX/XY chromosome system is highly conserved in therian mammals, dating back to just before the eutherian-marsupial split ∼180 Myr ago (Veyrunes et al. 2008). Given its age and apparently limited turnover, the mammal XX/XY system is often considered stable relative to sex chromosome systems in other vertebrate lineages (van Doorn and Kirkpatrick 2007; Veyrunes et al. 2008; Graves 2016). Nonetheless, variant sex chromosome systems have evolved in at least 20 eutherian mammal families (Saunders and Veyrunes 2021; Hughes et al. 2024). While much of the previous research on variant sex chromosome systems in mammals has focused on rodents (Fredga and Bulmer 1988; Marchal et al. 2003; Romanenko and Volobouev 2012; Saunders and Veyrunes 2021), variants are widespread in other mammalian clades including shrews (Sharman 1956), mongooses (Fredga 1965, 1972), and wallabies (Martin and Hayman 1966). The majority of these variants are neo-sex chromosomes, formed by one or more fusions — typically Robertsonian (Rb) translocations at the centromeres — of an autosome to a sex chromosome (Hughes et al. 2024). For example, the fusion of an autosome to a Y chromosome will produce an X_1_X_2_Y system, with the homolog of the fused autosome remaining as a second X. Such fusions lead to an odd number of chromosomes in one sex and are generally expected to be at least weakly deleterious (White et al. 1998; Ashley 2002; Barasc et al. 2011; Pennell et al. 2015). However, there are theoretical conditions under which neo-sex chromosomes may be favourable (Charlesworth and Charlesworth 1980; Charlesworth and Wall 1999), including linking pairs of locally adaptive alleles on separate chromosomes (Guerrero and Kirkpatrick 2014).

Multiple models have been proposed to describe the circumstances under which sex-autosome fusions are likely to become fixed, namely: sexual conflict, overdominance, and meiotic drive. Sexual conflict is the earliest theoretical framework for sex-autosome fusions (Fisher 1931; Charlesworth and Charlesworth 1980). For a locus that benefits one sex at a cost to the other, a sex-autosome fusion can resolve sexual conflict by restricting the locus to the sex it benefits (Rice 1984; van Doorn and Kirkpatrick 2007; Zhou and Bachtrog 2012; Matsumoto and Kitano 2016). Intralocus sexual conflict within a population may be impacted by environmental and ecological conditions (Berger et al. 2014; Dagilis et al. 2022). Specifically, sexual conflict may be more pronounced in populations in stable environments with few or mild seasonal changes relative to populations occupying novel or rapidly fluctuating environments (De Lisle et al. 2018; Plesnar-Bielak and Łukasiewicz 2021). In populations with pronounced levels of intralocus sexual conflict, the putative fitness benefits of resolving that conflict via a sex-autosome fusion could outweigh the expected costs of such rearrangements, increasing the likelihood of fixation (van Doorn and Kirkpatrick 2007; Matsumoto and Kitano 2016).

If, rather than being sexually antagonistic, an autosomal locus confers an advantage to heterozygotes (i.e. overdominance), then a sex-autosome fusion benefits the heterogametic sex (Charlesworth and Wall 1999). Whereas theory predicts that, in either of these scenarios, Y-autosome fusions will be more common than X-autosome fusions (Charlesworth and Charlesworth 1980; Charlesworth and Wall 1999), both types of fusion are approximately equally frequent in mammals (White 1973; Yoshida and Kitano 2012; Pennell et al. 2015; Hughes et al. 2024), suggesting the involvement of other processes.

Meiotic drive, the biased transmission of an allele causing non-Mendelian inheritance, is also predicted to shape the fixation of sex-autosome fusions. Meiotic drive *sensu stricto* occurs during female meiosis, where a driving locus — such as a centromere — can ensure the chromosome carrying it is preferentially transmitted to the oocyte rather than the polar body (Pardo-Manuel de Villena and Sapienza 2001*a*). Interestingly, whether the fused (metacentric) or unfused (acrocentric) chromosomes are preferentially transmitted in oogenesis differs between species (Pardo-Manuel de Villena and Sapienza 2001*a*). This difference in the ability of metacentric vs. acrocentric chromosomes to drive in oogenesis is termed meiotic polarity (Pardo-Manuel de Villena and Sapienza 2001*b*, 2001*a*; Yoshida and Kitano 2012; Blackmon et al. 2019). If meiotic polarity favours metacentrics then X-autosome fusions can readily become fixed whereas, if acrocentrics are favoured, Y-autosome fusions can accumulate because the Y chromosome never undergoes meiosis in oogenesis (Yoshida and Kitano 2012; Pokorná et al. 2014). As such, neo-sex chromosomes might be associated with different rates of chromosome evolution than XX/XY sex chromosomes, and with karyotypes that are biased towards metacentric or acrocentric chromosomes.

While meiotic drive is a compelling explanation for the emergence and fixation of neo-sex chromosomes and karyotype evolution in general, groups such as Carnivora exhibit rate variation in karyotype evolution that meiotic drive *sensu stricto* fails to explain (Jonika et al. 2024). When large climatic and environmental shifts reduce a species’ effective population size, fused chromosomes that typically exhibit heterozygote disadvantage (Lande 1979, 1985) can reach fixation by genetic drift (Lande 1985; Dobigny et al. 2005; Blackmon et al. 2024; Jonika et al. 2024; Mackintosh et al. 2024). An inverse correlation between species range size — a proxy for effective population size — and rates of chromosome fusion and fission in Carnivora suggests that genetic drift can strongly influence karyotype evolution (Jonika et al. 2024).

Clearly, a diverse set of phenomena may underlie the evolution and fixation of neo-sex chromosomes, and their exact contributions are likely to be taxon specific. Signals of sexual conflict are difficult to measure directly, but environmental stability has previously been associated with higher levels of sexual conflict (De Lisle et al. 2018). On the other hand, adaptive explanations should be reflected in ecological and environmental conditions that present opportunities to occupy a novel niche, which are generally associated with more stressful and cyclic environments where multiple strong selective pressures act (Hoffmann and Hercus 2000; Bell 2010; MacColl 2011). In such cases, extreme environmental conditions may enable the fixation of neo-sex chromosomes conferring a locally adaptive benefit. Alternatively, variable environments that create recurrent bottlenecks could result in the fixation of otherwise deleterious fusions via genetic drift. Finally, if sex-autosome fusions are primarily fixed by meiotic drive, with meiotic polarity favouring metacentric (or acrocentric) chromosomes, then karyotypes with neo-sex chromosomes should have a different proportion of metacentrics and acrocentrics than those without.

Here, we evaluated the potential contributions of drift, ecological selection, the environmental opportunity for sexual antagonism, and meiotic drive to the fixation of neo-sex chromosomes by testing for correlations between a suite of representative variables. Specifically, we compared species range size (an ecological proxy for effective population size), environmental conditions, and metacentric- or acrocentric-biased chromosome morphology in taxa with and without neo-sex chromosomes using phylogenetic logistic regression (Ives and Garland 2010), a modified *Mk* model of chromosome evolution (Blackmon et al. 2019, 2023), and stochastic mapping of sex-autosome fusion rates (Chien and Blackmon 2026). Three representative mammal families that encompass the diversity of sex-autosome fusions in Mammalia were included in our study: Herpestidae (mongooses, Y-autosome fusions), Soricidae (shrews, X-autosome fusions), and Phyllostomidae (New World leaf-nosed bats, both X-autosome fusions and combined X- and Y-autosome fusions).

## Methods

### Phylogenies

We selected three mammal families for analysis from the Upham et al. (2019) mammal phylogeny, each representing a different form of sex-autosome fusion. Only extant taxa were considered in our analyses. Ten species of herpestids (mongooses), out of 35 total, have two independent single fusions between the Y chromosome and an autosome (Fredga 1972; Murata et al. 2016), whereas ten species of soricids (shrews), out of 414 total, have a single fusion between the X chromosome and an autosome (Sharman 1956; Bulatova et al. 2019). In contrast, 29 species of phyllostomids (New World leaf-nosed bats), out of 205 total, exhibit X-autosome fusions, both individually and in combination with Y-autosome fusions (Baker et al. 1989; Noronha et al. 2010; Gomes et al. 2016) (Figure 1).

**Figure 1:**
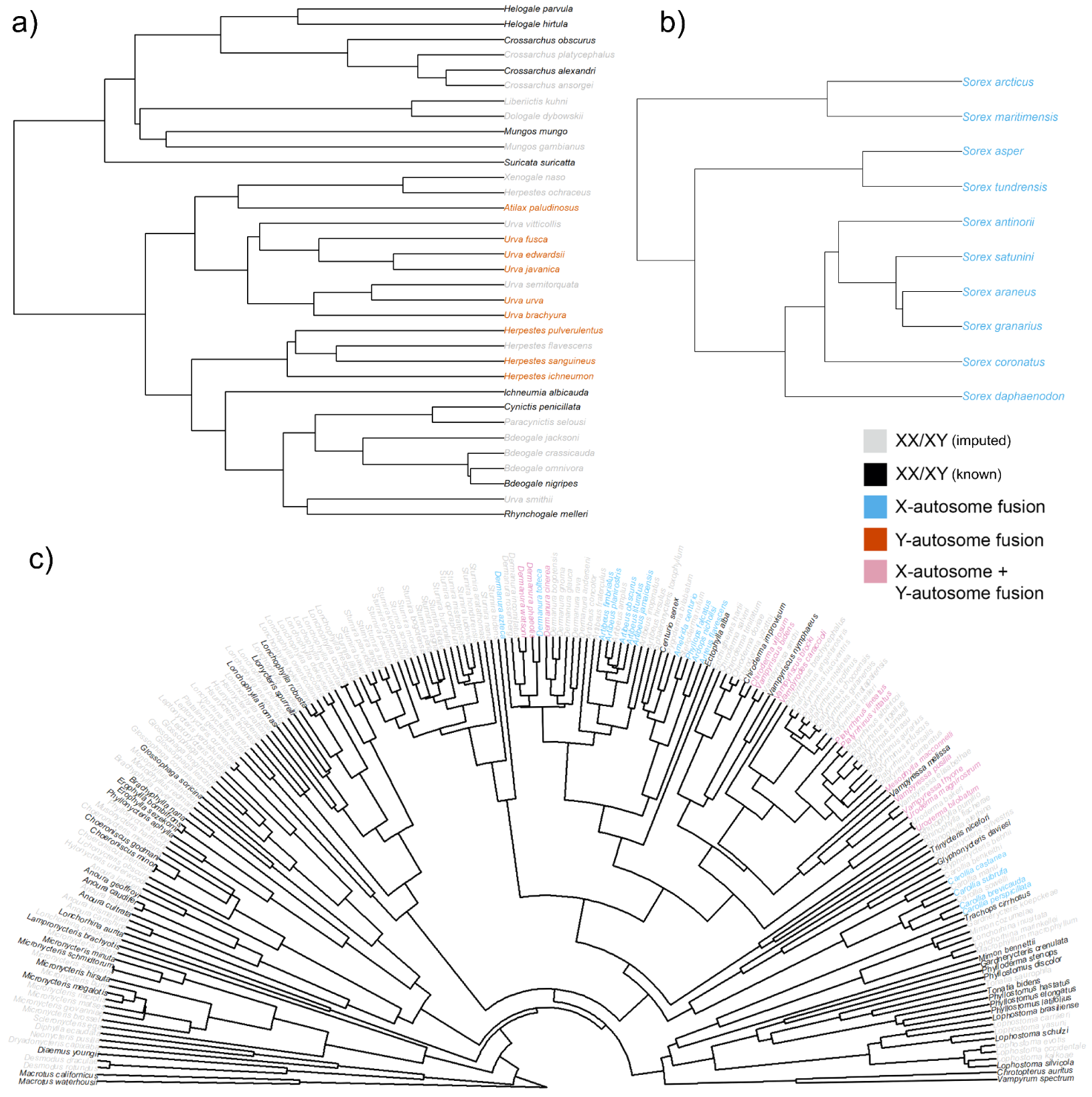
Phylogenetic trees of species in a) Herpestidae (9 variant species/16 with known karyotype/34 total), b) Soricidae (truncated to clade of 10 species containing known variant species only, excluding *Sorex kozlovi*/49 with known karyotype/413 total), and c) Phyllostomidae (29 variant species/71 with known karyotype/205 total), pruned from Upham et al.’s (2019) maximum clade credibility tree, with tips color-coded based on sex chromosome system. Species with X-autosome fusions are marked in blue, species with Y-autosome fusions are marked in orange, and species with both X- and Y-autosome fusions are marked in pink. Species known as XX/XY are marked in black, while those imputed as such are marked in gray.

For each of these three families, species names were reconciled with the Mammal Diversity Database (MDD) version 1.11 (Burgin et al. 2018) in both our datasets and the phylogenetic tree used in our analyses. We downloaded the maximum clade credibility consensus phylogeny, plus 1000 credible phylogenies from the posterior distribution of topologies, from Upham et al. (2019). Each tree was pruned to provide a separate family-level phylogeny for Herpestidae, Soricidae, and Phyllostomidae using phylogenetics tools from *ape* v5.8-1 (Paradis and Schliep 2019).

### Coding sex chromosome systems

Using the variant sex chromosome dataset from Hughes et al. (2024), each species was classified according to whether it had no sex-autosome fusion, an X-autosome fusion, a Y-autosome fusion, or both X- and Y-autosome fusions (Supplemental Table 1). When testing for correlations between presence of a sex-autosome fusion and our ecoclimatic variables (i.e. species range size alongside all climatic variables), we ran three sets of analyses. In the first (“XX/XY Default”) we assumed that species without karyotype data had no sex-autosome fusions (i.e. were XX/XY). In the second (“Parsimonious Karyotype”), where cytological information was unavailable for a species, we assigned the most parsimonious sex chromosome system, assuming the fewest possible transitions from XX/XY to variant systems. In Soricidae, where there is a single shared X-autosome fusion, the “XX/XY Default” and “Parsimonious Karyotype” datasets are identical. Finally, in the third set (“Known Karyotype”), we excluded all species without karyotype data. Phylogenies were visualised with tips color-coded according to sex chromosome system using plotting tools from *ape* v5.8-1 (Paradis and Schliep 2019) (Figure 1). Biogeographical maps of sex chromosome distributions were plotted using range data from TetrapodTraits v2.0.1 (Moura et al. 2024) and shapefile plotting functions in *ggplot2* (Wickham 2016) and *rnaturalearth* (Massicotte and South 2025) (Figure 2).

**Figure 2:**
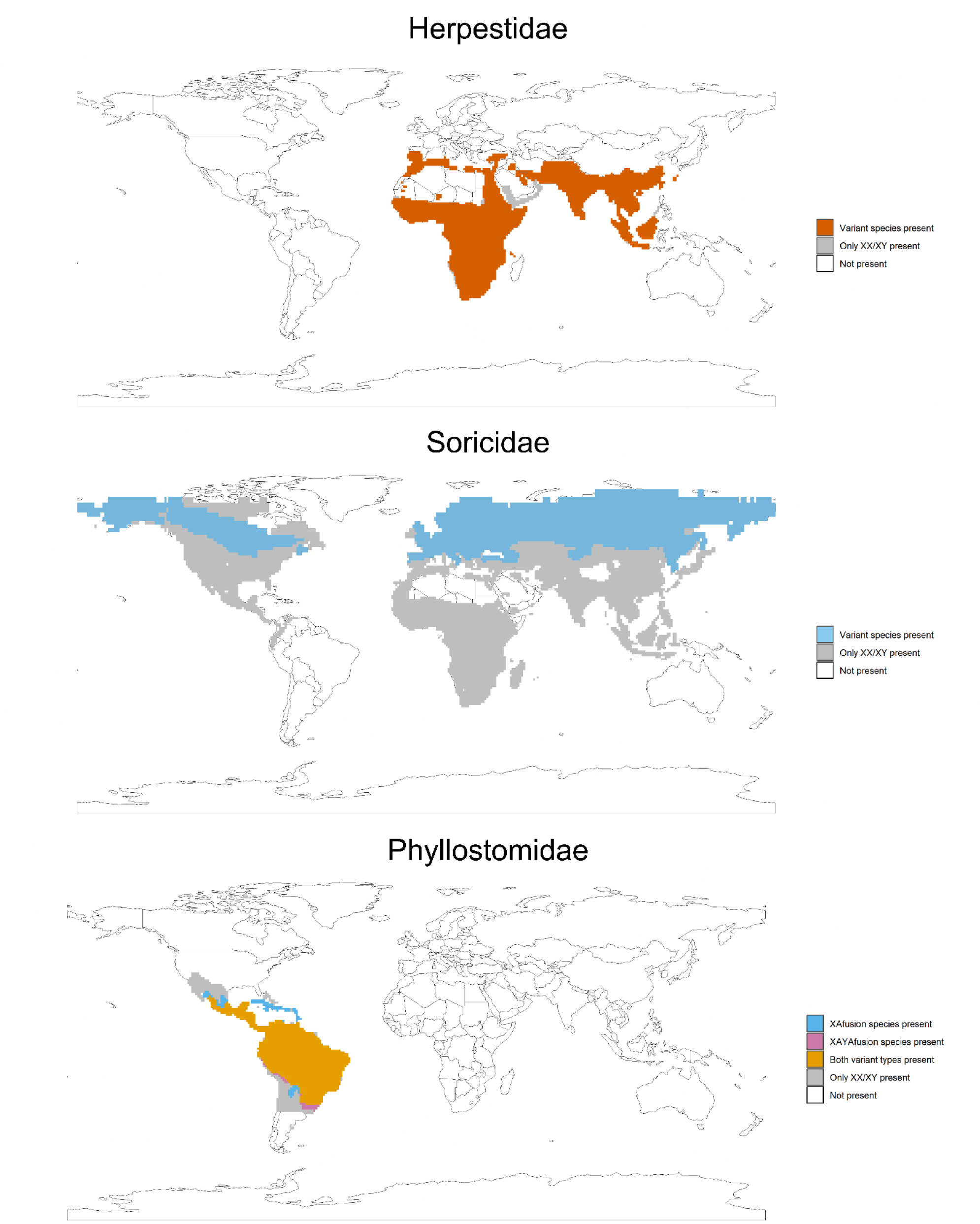
Distribution maps of all three mammal families studied. These maps are divided into 110km x 110km equal area grid cells, with species being grouped and the cells they occupy being colored based on their sex determination system. In Phyllostomidae, “both variant types” refers to areas where both species with X-autosome fusions and species with X- and Y-autosome fusions are present.

### Taxonomic uncertainty

Due to uncertainty in species delimitation in certain families, some tips were merged or removed from our analysis. In Herpestidae, *Urva auropunctata* (small Indian mongoose) was not included in the mammal phylogeny from Upham et al. (2019). We therefore considered it synonymous with *Urva javanica* (Javan mongoose), a species with a putatively identical karyotype that includes a shared Y-autosome fusion (Fredga 1972). This reduced the number of mongoose species to 34 total, and the number of variant mongoose species to 9. In Soricidae, *Sorex kozlovi* is traditionally placed in the *Sorex minutus* group (Bannikova et al. 2018). In the Upham et al. (2019) phylogeny, its position is instead imputed as sister to *S. daphaenodon* as the only assumed XX/XY species in a clade otherwise comprised of species with X-autosome fusions. Given the lack of available cytological evidence to support or falsify this likely spurious placement, *S. kozlovi* was excluded from our trees and analyses, leaving 413 species of shrew in total.

### Ecological and environmental variables

We obtained environmental data and range sizes for each of our taxa from TetrapodTraits v2.0.1 (Moura et al. 2024). We chose six environmental and climatic variables: latitude, annual mean temperature, temperature seasonality, annual precipitation, precipitation seasonality, and elevation. These six variables allowed us to test if neo-sex chromosome evolution is influenced by mean environmental conditions or their seasonal variation, whether directly (e.g. local adaptation) or indirectly (e.g. sexual conflict). We used the centroid of each species’ range for latitude and the within-range mean for the other five variables. Range size was included as an ecological proxy for effective population size, following the approach used by Jonika et al. (2024) to investigate rates of chromosome fusion and fission in Carnivora. Small populations in which the efficacy of selection is reduced might more readily fix deleterious chromosomal rearrangements via genetic drift.

### Chromosome morphology data

It has been argued that X-autosome fusions may become fixed by meiotic drive in mammals if metacentric chromosomes are preferentially transmitted over acrocentrics, and vice versa for Y-autosome fusions (Yoshida and Kitano 2012; Pokorná et al. 2014). In this case, the fixation of sex-autosome fusions could be a facet of karyotype-wide trends towards particular chromosome morphologies. To test this, we calculated the proportion of acrocentric chromosomes in the haploid female karyotype of each species, where available, from Baker et al. (1989), Blackmon et al. (2019), and from references cited in Table 1 of Hughes et al. (2024). For this same set of species, we derived the autosomal haploid chromosome count for each by subtracting 1 from the female haploid count to account for the X chromosome. Species without cytological data were excluded from the analyses where chromosome morphology variables were used as predictors.

**Table 1:**
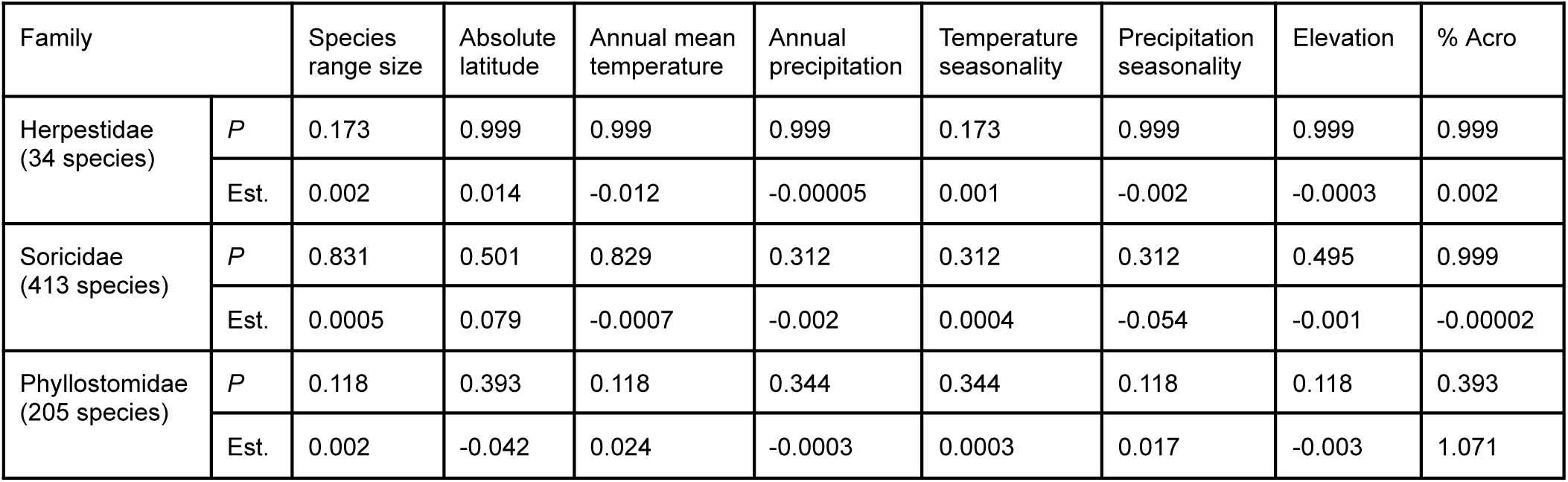
Phylogenetic logistic regression p-values (upper, corrected for false discovery rate) and estimates (lower) from correlation testing between seven ecoclimatic variables/percentage of acrocentric chromosomes and the presence of sex-autosome fusions in three mammal families using the consensus phylogeny. Results are shown for the “Assumed XX/XY” dataset except for tests of % acrocentric, which were only conducted using species in the “Known Karyotype” dataset.

### Hypothesis testing

All analyses were conducted in *R* v4.5.1. To assess whether environmental variables and species range size were correlated with sex-autosome fusions, we performed a phylogenetic logistic regression using the *phyloglm* function from the *phylolm* package v2.6.5 (Tung Ho and Ané 2014). We used the “logistic_MPLE” method and conducted 1000 independent bootstrap replicates using *phylolm*’s built-in bootstrap function. We treated the presence or absence of a sex-autosome fusion as our binary response variable, and fitted a separate *phyloglm* model for each of our variables. We initially used the maximum clade credibility consensus tree to calculate individual p-values, which were corrected for false discovery rate (FDR) using base R’s *p.adjust* function (Table 1).

To account for the possible influence of topological uncertainty on our models, we used the *tree_glm* function from the package *sensiPhy* v0.8.5 (Paterno et al. 2018). This allowed us to implement *phyloglm* across 1000 credible trees for all three mammal families of interest, and obtain a distribution of plausible parameter estimates (Table 2, Figure 3, Supplemental Figures 1—3). Both the consensus tree and the *sensiPhy* analyses were repeated on each of our three datasets (Tables 1—2, Supplemental Tables 2—5). For the “Known Karyotype” dataset, we dropped the tips of excluded species from all phylogenies. This left 16 species with known karyotype data out of 34 total in Herpestidae, 49 species with known karyotype out of 413 total in Soricidae, and 71 species with known karyotype data out of 205 in Phyllostomidae (Supplemental Table 1). In addition to the previous *phylolm* and *sensiPhy* analyses, we included new tests with the proportion of acrocentric chromosomes as the predictor variable (Tables 1—2, Supplemental Tables 2—5).

**Figure 3:**
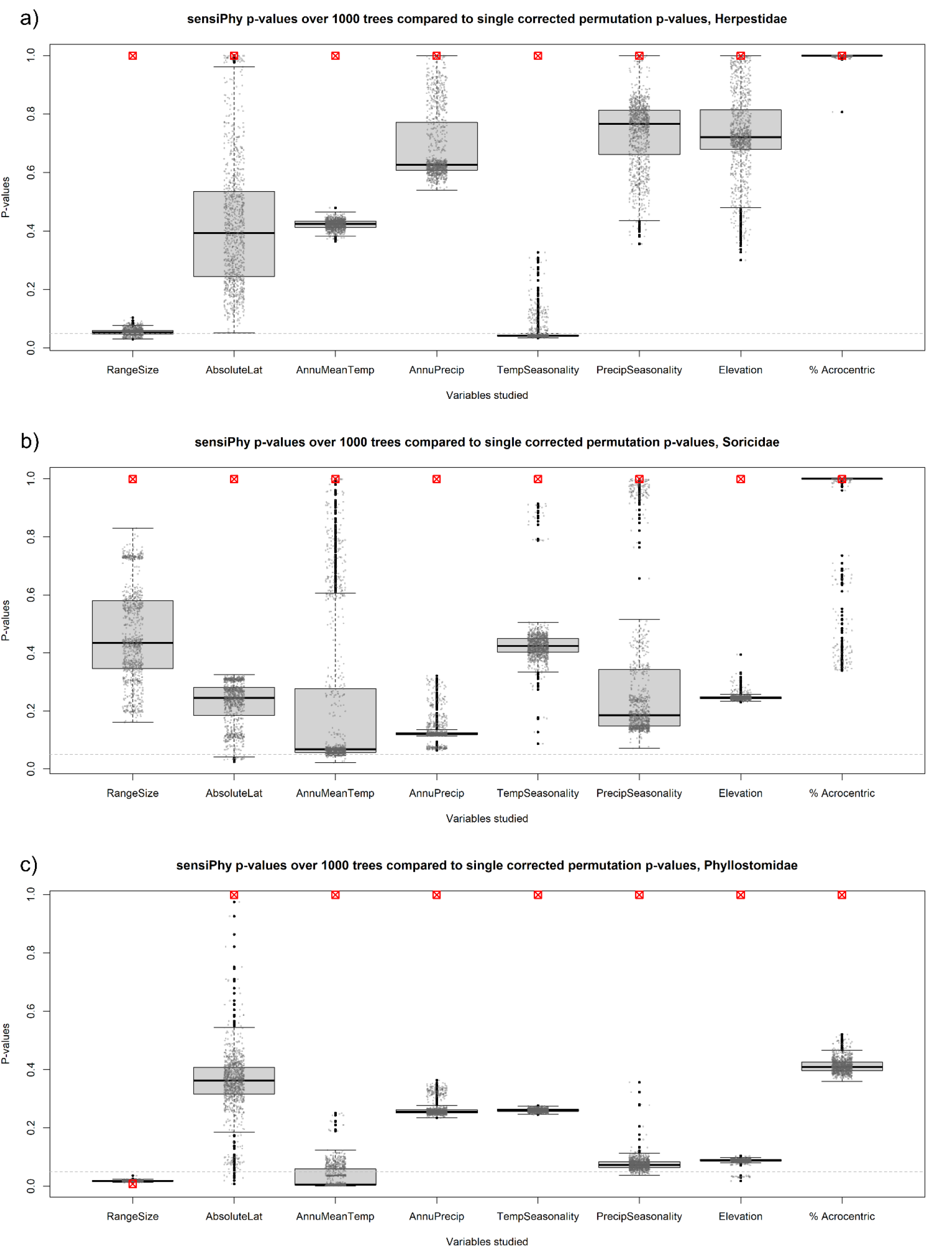
Box-and-whisker plots of p-values from 1000 hypothesis tests conducted over our variables of interest using *sensiPhy* for a) Herpestidae, b) Soricidae, and c) Phyllostomidae, the only family with an ecoclimatic variable exhibiting a corrected permutation p-value≤0.05 (species range size). Overlaid gray points are scatterplots of raw p-values from the same tests. Solid black points represent outlier data points in the box-and-whisker plots. Boxed red X’s indicate permutation p-values for each test calculated via the formula 1 − [(Σ(*number of tests wit*ℎ *pval*≤0. 05) + 1) ÷ (1000 + 1)] after false discovery rate correction. Gray dashed line indicates the p≤0.05 threshold. Tests of % Acrocentric were only conducted using species with known karyotypes.

**Table 2:** Mean p-values (upper) and estimates (center), and permutation p-values calculated via the formula 1 − [(Σ(*number of tests wit*ℎ *pval*≤0. 05) + 1) ÷ (1000 + 1)] (lower, corrected for false discovery rate) from *sensiPhy* analysis over 1000 credible trees for each family, for tested correlations between ecoclimatic variables/% acrocentric chromosomes and sex-autosome fusions. Statistically significant values (p≤0.05) in bold. Results are shown for the “Assumed XX/XY” dataset except for tests of % acrocentric, which were only conducted using species in the “Known Karyotype” dataset.

| Family |  | Species range size | Absolute latitude | Annual mean temperature | Annual precipitation | Temperature seasonality | Precipitation seasonality | Elevation | % Acro |
| --- | --- | --- | --- | --- | --- | --- | --- | --- | --- |
| Herpestidae (34 species) | Mean $p$ | 0.054 | 0.423 | 0.422 | 0.693 | 0.053 | 0.732 | 0.679 | 0.999 |
|  | Est. | 0.002 | 0.034 | -0.013 | 0.0002 | 0.0006 | 0.002 | -0.00006 | 0.002 |
|  | Permutation p-values | 0.999 | 0.999 | 0.999 | 0.999 | 0.999 | 0.999 | 0.999 | 0.999 |
| Soricidae (413 species) | Mean $p$ | 0.452 | 0.219 | 0.214 | 0.135 | 0.437 | 0.314 | 0.246 | 0.950 |
|  | Est. | 0.001 | 0.085 | -0.012 | -0.002 | 0.0002 | -0.044 | -0.0009 | -0.123 |
|  | Permutation p-values | 0.999 | 0.999 | 0.999 | 0.999 | 0.999 | 0.999 | 0.999 | 0.999 |
| Phyllostomidae (205 species) | Mean $p$ | **0.019** | 0.355 | **0.032** | 0.261 | 0.261 | 0.076 | 0.087 | 0.413 |
|  | Est. | 0.002 | -0.025 | 0.025 | -0.0004 | 0.0003 | 0.016 | -0.004 | 0.963 |
|  | Permutation p-values | **0.008** | 0.999 | 0.999 | 0.999 | 0.999 | 0.999 | 0.999 | 0.999 |

In some *sensiPhy* tests, we found that regression analyses were not being completed for the full set of 1000 phylogenies. In these cases, we gradually lowered the “btol” parameter from its default (50) until all 1000 trees were successfully tested. To test for significance of these results, we applied a permutation test p-value framework where we used the formula 1 − [(Σ(*number of tests wit*ℎ *pval*≤0. 05) + 1) ÷ (1000 + 1)] to derive a single p-value for each variable tested in *sensiPhy*, which was then corrected for FDR using base R’s *p.adjust* function (Table 2; Supplemental Table 3). In the sole case where all 1000 tests landed below the p-value ≤ 0.05 threshold, we set the uncorrected permutation p-value to 0.001.

For the taxa in the “Known Karyotype” dataset, we conducted two additional tests. First, we tested whether variant sex chromosomes were associated with changes in rates of chromosome fusion and fission across the whole karyotype (Figure 4). We used the *R* package *chromePlus* v2.0 (Blackmon et al. 2019, 2023) to specify a model of chromosome number evolution, with XY or variant sex chromosomes as our binary variable. For each of the two states, we aimed to infer the rates of fusions, fissions (chromosome number -1 and +1, respectively), and transitions between states. We fit this model to the set of 1000 credible trees using the *mcmc* function in *diversitree* v0.10-1 (FitzJohn 2012), running for 10,000 iterations each and discarding the first 1000 as burn-in. Following Blackmon et al. (2019), we calculated the mean rate difference at each iteration as the sum of the fusion and fission rates in state 2 (variant) divided by two, minus the sum of the fusion and fission rates in state 1 (XY) divided by two. Association between sex-autosome fusions and shifts in the rate of chromosome number evolution would suggest that sex-autosome fusions evolve and fix as part of karyotype-wide patterns of fusion and fission. This could include biases towards a particular chromosome morphology, as described by the meiotic polarity model (Pardo-Manuel de Villena and Sapienza 2001*a*; Yoshida and Kitano 2012; Blackmon et al. 2019). Finally, we also tested whether observed rates of sex-autosome fusion exceeded null expectations (i.e. all chromosomes are equally likely to undergo fusion) via stochastic mapping of estimated rates over the set of 1000 credible trees, with 100 simulations run for each tree. Following Chien and Blackmon (2026), we used an equation described by Anderson et al. (2020) to calculate the weighted sum of expected sex-autosome fusions, which considers the duration of each karyotype state across the inferred stochastic maps. This serves as the null expectation, which was then compared with the observed data (Supplemental Figure 4). Single-fusion events (e.g. X-A or Y-A fusion) were treated as separate karyotype states from double-fusion events (combined X-A + Y-A fusion).

**Figure 4:**
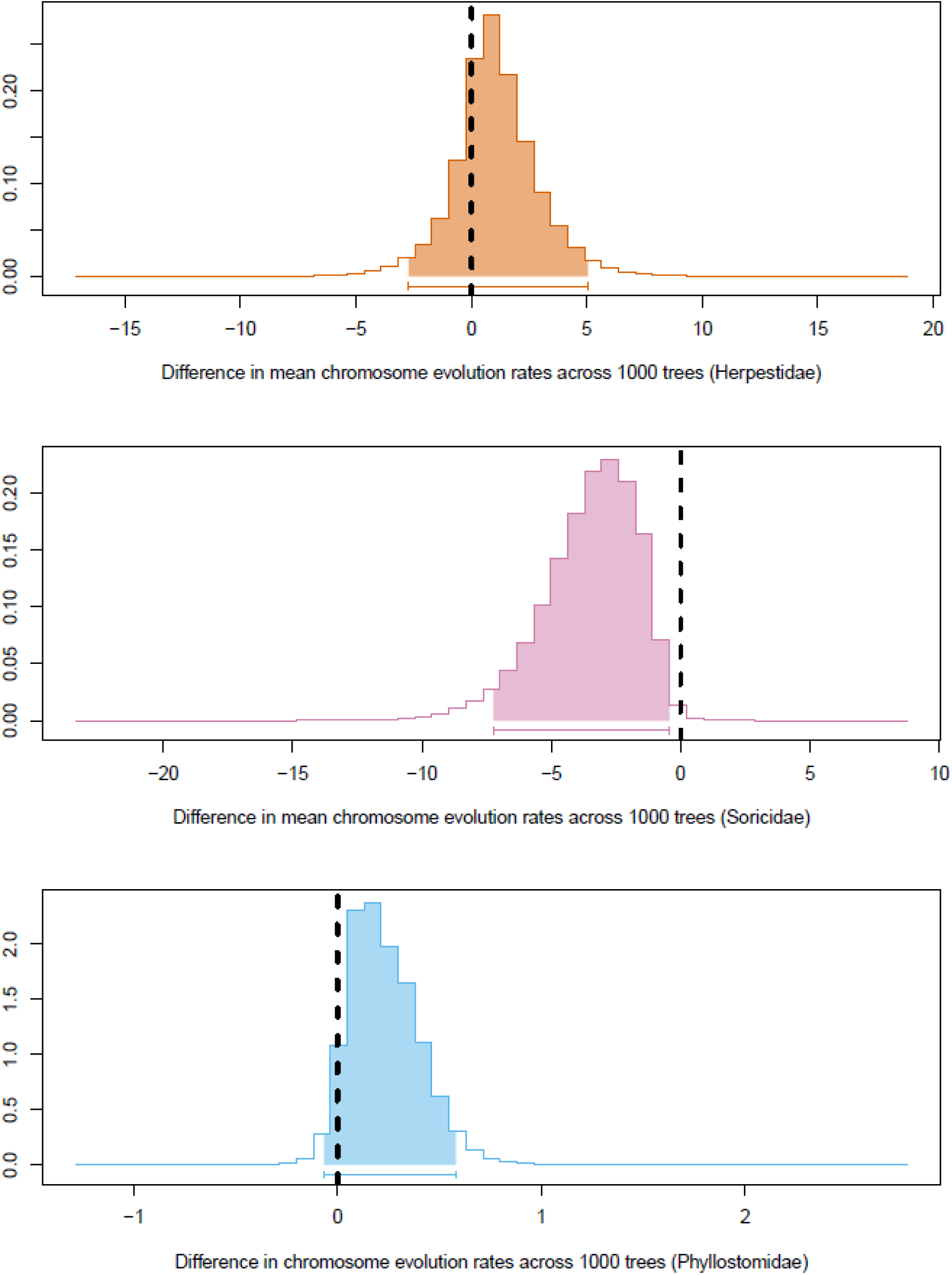
Differences in mean rates (per millions of years) of chromosome evolution between taxa with XX/XY and variant sex chromosomes in Herpestidae (upper), Soricidae (center), and Phyllostomidae (lower). The 95% credible interval is shown by the line beneath the probability distribution. Each shows the rates as inferred across 1000 credible trees.

## Results

Hypothesis testing using phylogenetic logistic regression was conducted for 652 total mammal species across three families, of which 48 species had known variant sex chromosome systems. In Herpestidae, we were able to analyze 34 total species, with 9 of these species having a Y-autosome fusion. In Soricidae, out of 413 total analyzed species, 10 have an X-autosome fusion. In Phyllostomidae, out of 205 total analyzed species, 15 possess an X-autosome fusion and 14 have both an X- and Y-autosome fusion, totalling 29 variant species. Hypothesis testing using karyotype data was conducted in 16/34 Herpestidae species, 49/413 Soricidae species, and 71/205 Phyllostomidae species. Under the “Parsimonious Karyotype” dataset, 5 additional herpestid species and 37 additional phyllostomid species were considered to have a variant sex chromosome system, totaling 14 variant herpestid species and 66 variant phyllostomid species overall (Supplemental Table 1). We assigned these species as “variant” or “XY” without consideration for the specific type of sex chromosome variant, as our logistic regression analysis only requires a binary input.

In analyses conducted with Upham et al.’s (2019) maximum clade credibility tree using the “logistic_MPLE” method with 1000 bootstrap replicates and FDR correction, no statistically significant relationship was detected between any of our selected ecoclimatic predictor variables and the presence of a sex-autosome fusion (Table 1, Supplemental Tables 2 and 4). There was also no statistically significant relationship between the proportion of acrocentric chromosomes in species’ karyotypes and the presence of a sex-autosome fusion (Table 1).

However, our *sensiPhy* analyses, which used the same predictor variables and 1000 credible phylogenies, demonstrated that associations between variant sex chromosomes and predictor variables are highly dependent on tree topology (Figure 3, Supplemental Figures 1—3, 5—6). Specifically, sex chromosome variants were significantly correlated with both species range size and annual mean temperature in Phyllostomidae when using the “Assumed XX/XY” dataset in the majority of the 1000 phylogenetic trees provided. Table 2 shows the mean estimates and permutation p-values after FDR correction for each variable over 1000 hypothesis tests in each family. With this dataset, under a limited number of phylogenetic hypotheses, there was also a significant correlation between neo-sex chromosomes and both range size and temperature seasonality in Herpestidae (Figure 3, Table 2).

When re-running *phylolm* and *sensiPhy* analyses in taxa with the “Known Karyotype” or “Parsimonious Karyotype” datasets, we found no evidence of correlation between ecoclimatic variables and sex-autosome fusions (Supplemental Table 2—5).

Our tests for an association between sex chromosome type and changes in the rate of chromosome evolution (i.e. fusions and fissions) yielded inconsistent results between families (Figure 4). For Herpestidae and Phyllostomidae, the 95% credible interval of the probability distribution overlapped zero, which suggests there is no difference in rates of chromosome evolution between XX/XY species and those with sex-autosome fusions in these families. In Soricidae, the credible interval is entirely negative. This may be evidence of lower rates of chromosome evolution in soricids with X-autosome fusions than those without. However, this result is more likely to be an artifact of the putative single event that led to the fixation of the X-autosome fusion in this specific clade of shrews. Our tests comparing observed sex-autosome fusion rates to null expectations recovered overlapping 95% credible intervals between the two probability distributions (Supplemental Figure 4), indicating no evidence for a difference in rates.

## Discussion

Whereas the influence of sexual and intragenomic conflict on the evolution of neo-sex chromosome systems has been studied extensively (van Doorn and Kirkpatrick 2007; Furman et al. 2020; Blackmon et al. 2024; Hughes et al. 2024; Minovic and Nozawa 2024), the potential influence of ecological and climatic factors on the fixation of neo-sex chromosomes is less understood. Here, we linked a comprehensive dataset of neo-sex chromosomes in three families of mammals with ecological and environmental data. We used phylogenetic logistic regressions to test whether effective population size (using range size as a proxy) and environmental associations can influence neo-sex chromosome evolution in mammals. We also tested if neo-sex chromosomes are associated with the proportion of acrocentric chromosomes in a karyotype and shifts in rates of chromosome fusion and fission. Results using a single consensus phylogeny indicate that the evolution of neo-sex chromosomes in mammals is likely not influenced by range size nor by the environmental variables we studied. Lack of association between environment and neo-sex chromosomes also rules out a role for sexual antagonism, which is expected to be a stronger force in stable relative to fluctuating environments (De Lisle et al. 2018; Plesnar-Bielak and Łukasiewicz 2021). Likewise, we found no relationship between autosome morphology and neo-sex chromosomes; in contrast to expectations from meiotic drive, patterns of rate shifts for chromosome fusions and fissions were inconsistent between taxa.

However, using a set of 1000 credible phylogenies, we found that a subset of these inferences are sensitive to tree topology. Using this set of 1000 trees, we found significant associations between neo-sex chromosomes and both range size and annual mean temperature in Phyllostomidae (Figure 3, Table 2). In Herpestidae, approximately one-third (35.8%) of topologies also yield a significant relationship between neo-sex chromosomes and range size, with this proportion increasing to more than three-quarters (83.5%) of topologies for temperature seasonality. No tested relationships were statistically significant when restricted to species with known karyotypes, reinforcing the importance of collecting cytological data for all species.

Previous literature has linked range size and temperature with patterns of genome evolution. For example, polyploidy in multiple animal taxa is associated with latitude and temperature seasonality (David 2022). In Carnivora, species with smaller range sizes show elevated rates of chromosome fusion and fission (Jonika et al. 2024). Results for Herpestidae, the sole carnivoran family in our dataset, suggest the opposite pattern, with a significant but very weakly positive relationship between range size and sex-autosome fusions under some (358/1000) phylogenetic hypotheses (Figure 3, Table 2). However, we found no difference in overall rates of chromosome evolution for Herpestidae with XX/XY or Y-autosome fusions (Figure 4). This may reflect the extremely stable karyotypes of most carnivorans (excepting Ursidae and Canidae) (Perelman et al. 2004; Nie et al. 2012) and is consistent with the observation that meiotic drive alone cannot explain patterns of karyotype morphology in Carnivora (Blackmon et al. 2019; Jonika et al. 2024). Finally, evidence for a significant but very weakly positive relationship between temperature seasonality and sex-autosome fusions under a majority of phylogenetic hypotheses in Herpestidae (Figure 3a) motivates future analyses using more exact measures of thermal clines.

Using the “Assumed XX/XY” dataset, Phyllostomidae also showed a significant but weakly positive relationship between range size and neo-sex chromosomes in *sensiPhy* (Figure 3, Table 2) that does not appear in our single-tree analysis. The direction of this correlation in both herpestids and phyllostomids is opposed to that predicted if drift in small populations drives the fixation of sex-autosome fusions. Given that range size is an imperfect proxy for effective population size, it would be of interest to repeat these analyses using genetic estimates of effective population size. However, it is also possible that sex-autosome fusions confer a locally adaptive benefit to variant species. Additional taxonomic and genomic comparisons are necessary to tease apart these possibilities.

Like Herpestidae, phyllostomids with neo-sex chromosomes do not differ in rates of chromosome fusion and fission compared to those with XX/XY sex chromosomes (Figure 4), nor in the proportion of acrocentric chromosomes. Observed rates of sex-autosome fusion within these two families also overlapped with expectations under stochastic mapping, supporting a lack of difference in rates between species with and without variant sex chromosomes (Supplemental Figure 4). We note two caveats to these results. First, because our *chromePlus* analysis required a binary categorization of variant or non-variant sex chromosomes, the analysis failed to capture the complex patterns of independent X-autosome fusions and subsequent independent additions of Y-autosome fusions in Phyllostomidae, despite neo-X chromosomes being associated with metacentrics and neo-Y chromosomes with acrocentrics (Yoshida and Kitano 2012). Second, our test for different rates of karyotype evolution in species with and without sex-autosome fusions did not evaluate whether some clades were more likely to fix sex-autosome fusions than others. This could be important given switches in meiotic polarity across lineages (Blackmon et al. 2019), and could be tested with finer-scale cladewise comparisons.

In Soricidae, for which we found no evidence of a relationship between neo-sex chromosomes and any of our tested variables, species with X-autosome fusions were inferred to have lower rates of chromosome evolution that those with XX/XY sex chromosomes. This may reflect the fact that the small number of shrews with X-autosome fusions form a clade (Figure 1b) with a small range of chromosome numbers (autosomal haploid numbers of 10-15), relative to the entire family (autosomal haploid numbers of 10-32). As such, this clade of shrews likely experienced multiple autosome-autosome fusions, and now show limited chromosome number variation between species. Thus, observed evolutionary rate differences in Soricidae may be an artifact of reduced chromosome numbers within the single clade with X-autosome fusions.

A limitation of our approach is that species range sizes and environmental variables from the TetrapodTraits dataset provide a current representation of species’ biogeography and therefore may not capture past conditions these taxa experienced. This may be important for taxa whose ecology and population structure were shaped by paleoclimatic conditions distinct from those in their contemporary range. For example, some West African gerbils in the genus *Taterillus* have X-autosome fusions and marked differences in chromosome numbers. It has been suggested that these differences in karyotypes were fixed in small populations between cycles of Saharan expansion and contraction, with rivers limiting dispersal during periods of contraction (Dobigny et al. 2005). Incorporating models of historical shifts in ecology and demography would improve our framework.

As demonstrated by the *sensiPhy* analysis, tree topology can change the results of comparative analyses even when the same data are used. The mammal tree from Upham et al. (2019) used in our initial phylogenetic logistic regression with *phylolm* is a maximum clade credibility tree, and inherently represents a single phylogenetic hypothesis. The use of only one tree in phylogenetic comparative methods has been identified as a potential source of statistical bias, as this approach fails to account for phylogenetic uncertainty and the possibility of different evolutionary hypotheses (Donoghue and Ackerly 1996; Hernández et al. 2013; Rangel et al. 2015). Incorporation of multiple phylogenies provides a more informative measure of uncertainty in comparative analysis than can be achieved using a single tree (Figure 3, Supplemental Figures 5—6). We also note that the only *sensiPhy* tests where a majority of the correlations were significant were those using the “Assumed XX/XY” dataset, reaffirming the need to consider multiple evolutionary hypotheses.

In summary, for three families of mammals, we found no evidence for strong of consistent associations between neo-sex chromosomes and environmental conditions or geographic range size. We also found no clear evidence for taxon-specific differences in rates of chromosome evolution between species with and without neo-sex chromosomes. Limitations of sample size and data resolution notwithstanding, our findings suggest that other conditions and processes are more important drivers of neo-sex chromosome evolution and fixation in mammals. Importantly, the results of this study highlight the value of including multiple phylogenetic hypotheses in comparative analysis.

## Acknowledgements

We thank Louise Heitzmann, Daniel Moen, and Nathan Upham for their advice regarding analyses and statistical testing. We also thank Heath Blackmon and Gustavo Paterno for their assistance in using *chromePlus* and *sensiPhy*, respectively.

## Statement of Authorship

J.J.H. conceived the work; all authors designed the analyses; J.C. wrote code for and performed the analyses; J.J.H. and J.C. drafted the original manuscript; all authors reviewed and edited the manuscript; J.J.H. and P.C. supervised the research.

## Data and Code Availability

All data and code are stored on Github (JimmyKChoi/Variant-Sex-Chromosome-Systems-Bioclimatic-Variables), and will be archived at Zenodo upon acceptance.

## References

Ashley, T. 2002. X-Autosome translocations, meiotic synapsis, chromosome evolution and speciation. Cytogenetic and Genome Research 96:33–39.

Baker, R. J., C. S. Hood, and R. L. Honeycutt. 1989. Phylogenetic relationships and classification of the higher categories of the New World bat family Phyllostomidae. Systematic Biology 38:228–238.

Bannikova, A. A., D. Chernetskaya, A. Raspopova, D. Alexandrov, Y. Fang, N. Dokuchaev, B. Sheftel, et al. 2018. Evolutionary history of the genus *Sorex* (Soricidae, Eulipotyphla) as inferred from multigene data. Zoologica Scripta 47:518–538.

Barasc, H., N. Mary, R. Letron, A. Calgaro, A. M. Dudez, N. Bonnet, Y. Lahbib-Mansais, et al. 2011. Y-autosome translocation interferes with meiotic sex inactivation and expression of autosomal genes: a case study in the pig. Sexual Development 6:143–150.

Bell, G. 2010. Fluctuating selection: the perpetual renewal of adaptation in variable environments. Philosophical Transactions of the Royal Society B: Biological Sciences 365:87–97.

Berger, D., K. Grieshop, M. I. Lind, J. Goenaga, A. A. Maklakov, and G. Arnqvist. 2014. INTRALOCUS SEXUAL CONFLICT AND ENVIRONMENTAL STRESS. Evolution 68:2184–2196.

Blackmon, H., M. Chin, and M. Jonika. 2023. coleoguy/chromePlus: chromePlus. Zenodo.

Blackmon, H., M. M. Jonika, J. M. Alfieri, L. Fardoun, and J. P. Demuth. 2024. Drift drives the evolution of chromosome number I: The impact of trait transitions on genome evolution in Coleoptera. Journal of Heredity 115:173–182.

Blackmon, H., J. Justison, I. Mayrose, and E. E. Goldberg. 2019. Meiotic drive shapes rates of karyotype evolution in mammals. Evolution 73:511–523.

Britton-Davidian, J., J. Catalan, M. da Graça Ramalhinho, G. Ganem, J.-C. Auffray, R. Capela, M. Biscoito, et al. 2000. Rapid chromosomal evolution in island mice. Nature 403:158–158.

Bulatova, N. S., L. S. Biltueva, S. V. Pavlova, N. S. Zhdanova, and J. Zima. 2019. Chromosomal differentiation in the common shrew and related species. Pages 134–185 in J. Zima, J. B. Searle, and P. D. Polly, eds. Shrews, chromosomes and speciation, Cambridge studies in morphology and molecules: new paradigms in evolutionary biology. Cambridge University Press, Cambridge.

Burgin, C. J., J. P. Colella, P. L. Kahn, and N. S. Upham. 2018. How many species of mammals are there? Journal of Mammalogy 99:1–14.

Charlesworth, B., and J. D. Wall. 1999. Inbreeding, heterozygote advantage and the evolution of neo-X and neo-Y sex chromosomes. Proceedings of the Royal Society B: Biological Sciences 266:51.

Charlesworth, D., and B. Charlesworth. 1980. Sex differences in fitness and selection for centric fusions between sex-chromosomes and autosomes. Genetics Research 35:205–214.

Chesser, R. K., and R. J. Baker. 1986. On Factors Affecting the Fixation of Chromosomal Rearrangements and Neutral Genes: Computer Simulations. Evolution 40:625–632.

Chien, S., and H. Blackmon. 2026. Chromosomal rearrangements: tempo and mode of karyotype evolution in Scarabaeoidea. Journal of Evolutionary Biology 39:699–708.

Dagilis, A. J., J. M. Sardell, M. P. Josephson, Y. Su, M. Kirkpatrick, and C. L. Peichel. 2022. Searching for signatures of sexually antagonistic selection on stickleback sex chromosomes. Philosophical Transactions of the Royal Society B: Biological Sciences 377:20210205.

David, K. T. 2022. Global gradients in the distribution of animal polyploids. Proceedings of the National Academy of Sciences 119:e2214070119.

De Lisle, S. P., D. Goedert, A. M. Reedy, and E. I. Svensson. 2018. Climatic factors and species range position predict sexually antagonistic selection across taxa. Philosophical Transactions of the Royal Society B: Biological Sciences 373:20170415.

Dobigny, G., V. Aniskin, L. Granjon, R. Cornette, and V. Volobouev. 2005. Recent radiation in West African *Taterillus* (Rodentia, Gerbillinae): the concerted role of chromosome and climatic changes. Heredity 95:358–368.

Donoghue, M. J., and D. D. Ackerly. 1996. Phylogenetic uncertainties and sensitivity analyses in comparative biology. Philosophical Transactions of the Royal Society of London. Series B: Biological Sciences 351:1241–1249.

Fisher, R. A. 1931. The evolution of dominance. Biological Reviews 6:345–368.

FitzJohn, R. G. 2012. Diversitree: comparative phylogenetic analyses of diversification in R. Methods in Ecology and Evolution 3:1084–1092.

Fredga, K. 1965. New sex determining mechanism in a mammal. Nature 206:1176–1176.

Fredga, K. 1972. Comparative chromosome studies in mongooses (Carnivora, Viverridae). Hereditas 71:1–74.

Fredga, K., and M. G. Bulmer. 1988. Aberrant chromosomal sex-determining mechanisms in mammals, with special reference to species with XY females. Philosophical Transactions of the Royal Society of London. Series B, Biological Sciences 322:83–95.

Furman, B. L. S., D. C. H. Metzger, I. Darolti, A. E. Wright, B. A. Sandkam, P. Almeida, J. J. Shu, et al. 2020. Sex chromosome evolution: so many exceptions to the rules. Genome Biology and Evolution 12:750–763.

Gomes, A. J. B., C. Y. Nagamachi, L. R. R. Rodrigues, T. C. M. Benathar, T. F. A. Ribas, P. C. M. O’Brien, F. Yang, et al. 2016. Chromosomal phylogeny of Vampyressine bats (Chiroptera, Phyllostomidae) with description of two new sex chromosome systems. BMC Evolutionary Biology 16:119.

Graves, J. A. M. 2016. Did sex chromosome turnover promote divergence of the major mammal groups? BioEssays 38:734–743.

Guerrero, R. F., and M. Kirkpatrick. 2014. Local adaptation and the evolution of chromosome fusions. Evolution 68:2747–2756.

Hernández, C. E., E. Rodríguez-Serrano, J. Avaria-Llautureo, O. Inostroza-Michael, B. Morales-Pallero, D. Boric-Bargetto, C. B. Canales-Aguirre, et al. 2013. Using phylogenetic information and the comparative method to evaluate hypotheses in macroecology. (L. Harmon, ed.)Methods in Ecology and Evolution 4:401–415.

Hoffmann, A. A., and M. J. Hercus. 2000. Environmental Stress as an Evolutionary Force. BioScience 50:217.

Hughes, J. J., G. Lagunas-Robles, and P. Campbell. 2024. The role of conflict in the formation and maintenance of variant sex chromosome systems in mammals. Journal of Heredity 115:601–624.

Ives, A. R., and T. Garland. 2010. Phylogenetic logistic regression for binary dependent variables. Systematic Biology 59:9–26.

Jonika, M. M., K. T. Wilhoit, M. Chin, A. Arekere, and H. Blackmon. 2024. Drift drives the evolution of chromosome number II: The impact of range size on genome evolution in Carnivora. Journal of Heredity 115:524–531.

Lande, R. 1979. Effective deme sizes during long-term evolution estimated from rates of chromosomal rearrangement. Evolution 33:234–251.

Lande, R. 1985. The fixation of chromosomal rearrangements in a subdivided population with local extinction and colonization. Heredity 54:323–332.

MacColl, A. D. C. 2011. The ecological causes of evolution. Trends in Ecology & Evolution 26:514–522.

Mackintosh, A., R. Vila, S. H. Martin, D. Setter, and K. Lohse. 2024. Do chromosome rearrangements fix by genetic drift or natural selection? Insights from Brenthis butterflies. Molecular Ecology 33:e17146.

Marchal, J. A., M. J. Acosta, M. Bullejos, R. Díaz de la Guardia, and A. Sánchez. 2003. Sex chromosomes, sex determination, and sex-linked sequences in Microtidae. Cytogenetic and Genome Research 101:266–273.

Martin, P. G., and D. L. Hayman. 1966. A complex sex-chromosome system in the hare-wallaby *Lagorchestes conspicillatus* Gould. Chromosoma 19:159–175.

Massicotte, P., and A. South. 2025. rnaturalearth: World Map Data from Natural Earth.

Matsumoto, T., and J. Kitano. 2016. The intricate relationship between sexually antagonistic selection and the evolution of sex chromosome fusions. Journal of Theoretical Biology 404:97–108.

McAllister, B. F., S. L. Sheeley, P. A. Mena, A. L. Evans, and C. Schlötterer. 2008. Clinal distribution of a chromosomal rearrangement: a precursor to chromosomal speciation? Evolution 62:1852–1865.

Meisel, R. P. 2022. Ecology and the evolution of sex chromosomes. Journal of Evolutionary Biology 35:1601–1618.

Minovic, A., and M. Nozawa. 2024. Evolution of sex-biased genes in Drosophila species with neo-sex chromosomes: Potential contribution to reducing the sexual conflict. Ecology and Evolution 14:e11701.

Moura, M. R., K. Ceron, J. J. M. Guedes, R. Chen-Zhao, Y. V. Sica, J. Hart, W. Dorman, et al. 2024. A phylogeny-informed characterisation of global tetrapod traits addresses data gaps and biases. PLOS Biology 22:e3002658.

Murata, C., H. Sawaya, K. Nakata, F. Yamada, I. Imoto, and A. Kuroiwa. 2016. The cryptic Y-autosome translocation in the small Indian mongoose, *Herpestes auropunctatus*, revealed by molecular cytogenetic approaches. Chromosoma 125:807–815.

Nie, W., J. Wang, W. Su, D. Wang, A. Tanomtong, P. L. Perelman, A. S. Graphodatsky, et al. 2012. Chromosomal rearrangements and karyotype evolution in carnivores revealed by chromosome painting. Heredity 108:17–27.

Noronha, R. C. R., C. Y. Nagamachi, P. C. M. O’Brien, M. A. Ferguson-Smith, and J. C. Pieczarka. 2010. Meiotic analysis of XX/XY and neo-XX/XY sex chromosomes in Phyllostomidae by cross-species chromosome painting revealing a common chromosome 15-XY rearrangement in Stenodermatinae. Chromosome Research 18:667–676.

Paradis, E., and K. Schliep. 2019. ape 5.0: an environment for modern phylogenetics and evolutionary analyses in R. (R. Schwartz, ed.)Bioinformatics 35:526–528.

Pardo-Manuel de Villena, F., and C. Sapienza. 2001a. Female meiosis drives karyotypic evolution in mammals. Genetics 159:1179–1189.

Pardo-Manuel de Villena, F., and C. Sapienza. 2001b. Nonrandom segregation during meiosis: the unfairness of females. Mammalian Genome 12:331–339.

Paterno, G. B., C. Penone, and G. D. A. Werner. 2018. sensiPhy: An r-package for sensitivity analysis in phylogenetic comparative methods. Methods in Ecology and Evolution 9:1461–1467.

Pennell, M. W., M. Kirkpatrick, S. P. Otto, J. C. Vamosi, C. L. Peichel, N. Valenzuela, and J. Kitano. 2015. Y fuse? Sex chromosome fusions in fishes and reptiles. PLOS Genetics 11:e1005237.

Perelman, P. L., A. S. Graphodatsky, N. A. Serdukova, W. Nie, E. Z. Alkalaeva, B. Fu, T. J. Robinson, et al. 2004. Karyotypic conservatism in the suborder Feliformia (Order Carnivora). Cytogenetic and Genome Research 108:348–354.

Piálek, J., H. C. Hauffe, and J. B. Searle. 2005. Chromosomal variation in the house mouse. Biological Journal of the Linnean Society 84:535–563.

Plesnar-Bielak, A., and A. Łukasiewicz. 2021. Sexual conflict in a changing environment. Biological Reviews 96:1854–1867.

Pokorná, M., M. Altmanová, and L. Kratochvíl. 2014. Multiple sex chromosomes in the light of female meiotic drive in amniote vertebrates. Chromosome Research 22:35–44.

Rangel, T. F., R. K. Colwell, G. R. Graves, K. Fučíková, C. Rahbek, and J. A. F. Diniz-Filho. 2015. Phylogenetic uncertainty revisited: Implications for ecological analyses. Evolution 69:1301–1312.

Rice, W. R. 1984. Sex chromosomes and the evolution of sexual dimorphism. Evolution 38:735.

Romanenko, S. A., and V. Volobouev. 2012. Non-Sciuromorph Rodent Karyotypes in Evolution. Cytogenetic and Genome Research 137:233–245.

Saunders, P. A., and F. Veyrunes. 2021. Unusual mammalian sex determination systems: A cabinet of curiosities. Genes 12:1770.

Sharman, G. B. 1956. Chromosomes of the Common Shrew. Nature 177:941–942.

Tung Ho, L. si, and C. Ané. 2014. A linear-time algorithm for Gaussian and non-Gaussian trait evolution models. Systematic Biology 63:397–408.

Upham, N. S., J. A. Esselstyn, and W. Jetz. 2019. Inferring the mammal tree: Species-level sets of phylogenies for questions in ecology, evolution, and conservation. PLOS Biology 17:e3000494.

van Doorn, G. S., and M. Kirkpatrick. 2007. Turnover of sex chromosomes induced by sexual conflict. Nature 449:909–912.

Veyrunes, F., P. D. Waters, P. Miethke, W. Rens, D. McMillan, A. E. Alsop, F. Grützner, et al. 2008. Bird-like sex chromosomes of platypus imply recent origin of mammal sex chromosomes. Genome Research 18:965–973.

White, M. J. D. 1973. Animal cytology and evolution (Third Edition.). Cambridge University Press, Cambridge.

White, W. M., H. F. Willard, D. L. V. Dyke, and D. J. Wolff. 1998. The spreading of X inactivation into autosomal material of an X;autosome translocation: evidence for a difference between autosomal and X-chromosomal DNA. The American Journal of Human Genetics 63:20–28.

Wickham, H. 2016. ggplot2: elegant graphics for data analysis. Use R! (Second edition.). Springer international publishing, Cham.

Yin, Y., H. Fan, B. Zhou, Y. Hu, G. Fan, J. Wang, F. Zhou, et al. 2021. Molecular mechanisms and topological consequences of drastic chromosomal rearrangements of muntjac deer. Nature Communications 12:6858.

Yoshida, K., and J. Kitano. 2012. The contribution of female meiotic drive to the evolution of neo-sex chromosomes. Evolution 66:3198–3208.

Zhou, Q., and D. Bachtrog. 2012. Sex-specific adaptation drives early sex chromosome evolution in *Drosophila*. Science 337:341–345.

