## Supplementary Figure for "Ecoclimatic variables and karyotype features are poor predictors of neo-sex chromosome evolution in mammals"

**Supplemental Materials**

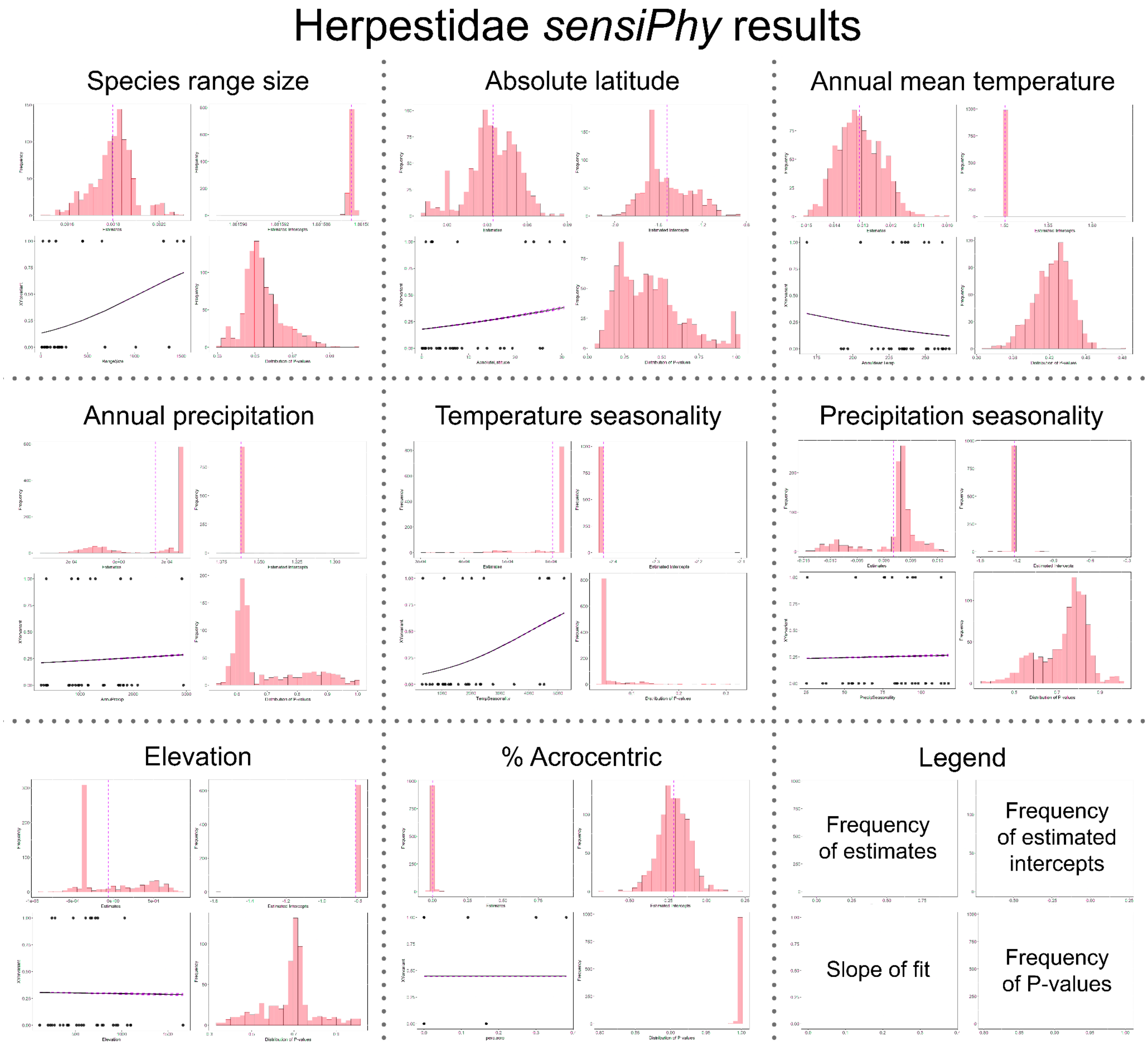

### Soricidae *sensiPhy* results

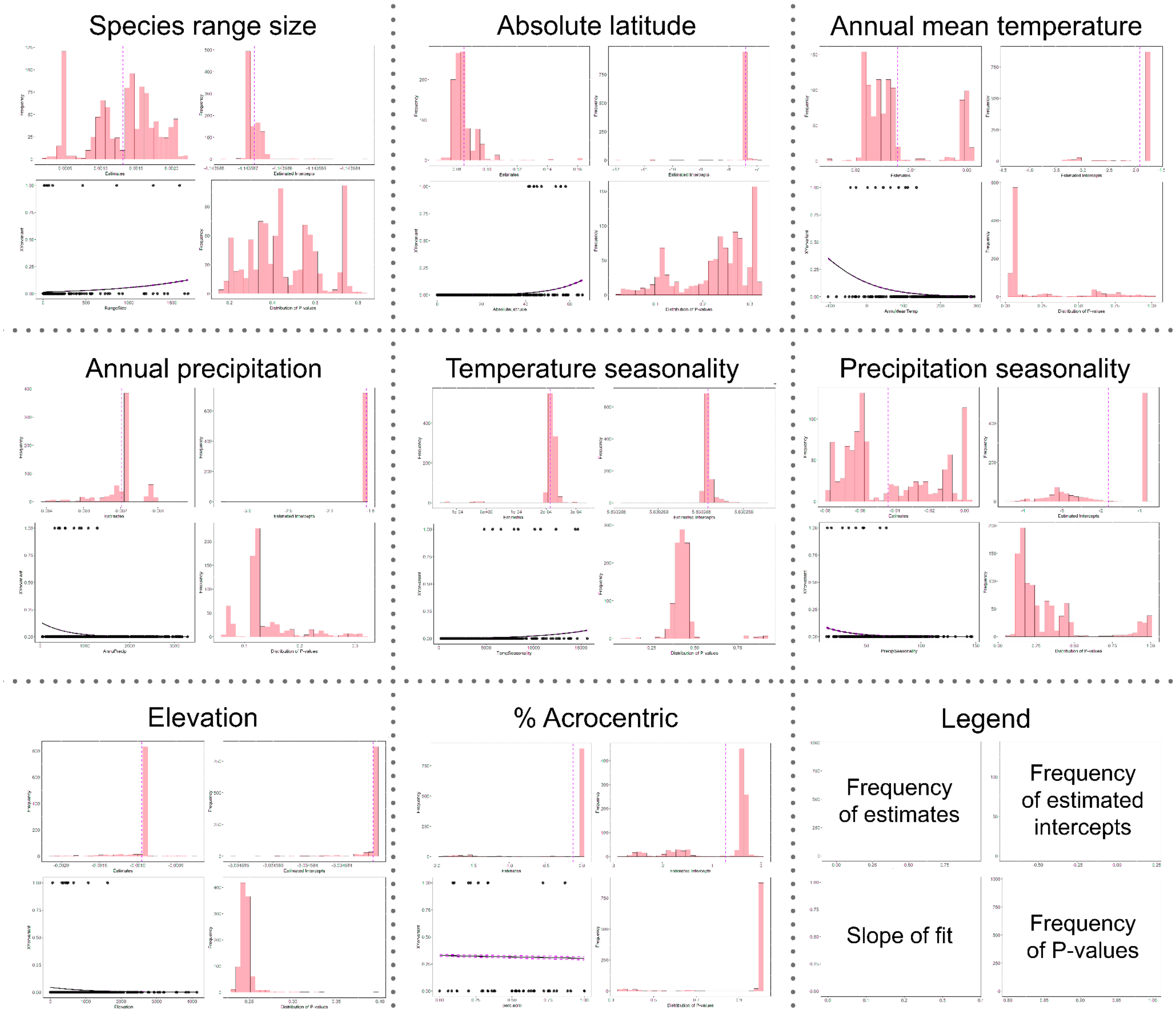

### Phyllostomidae *sensiPhy* results

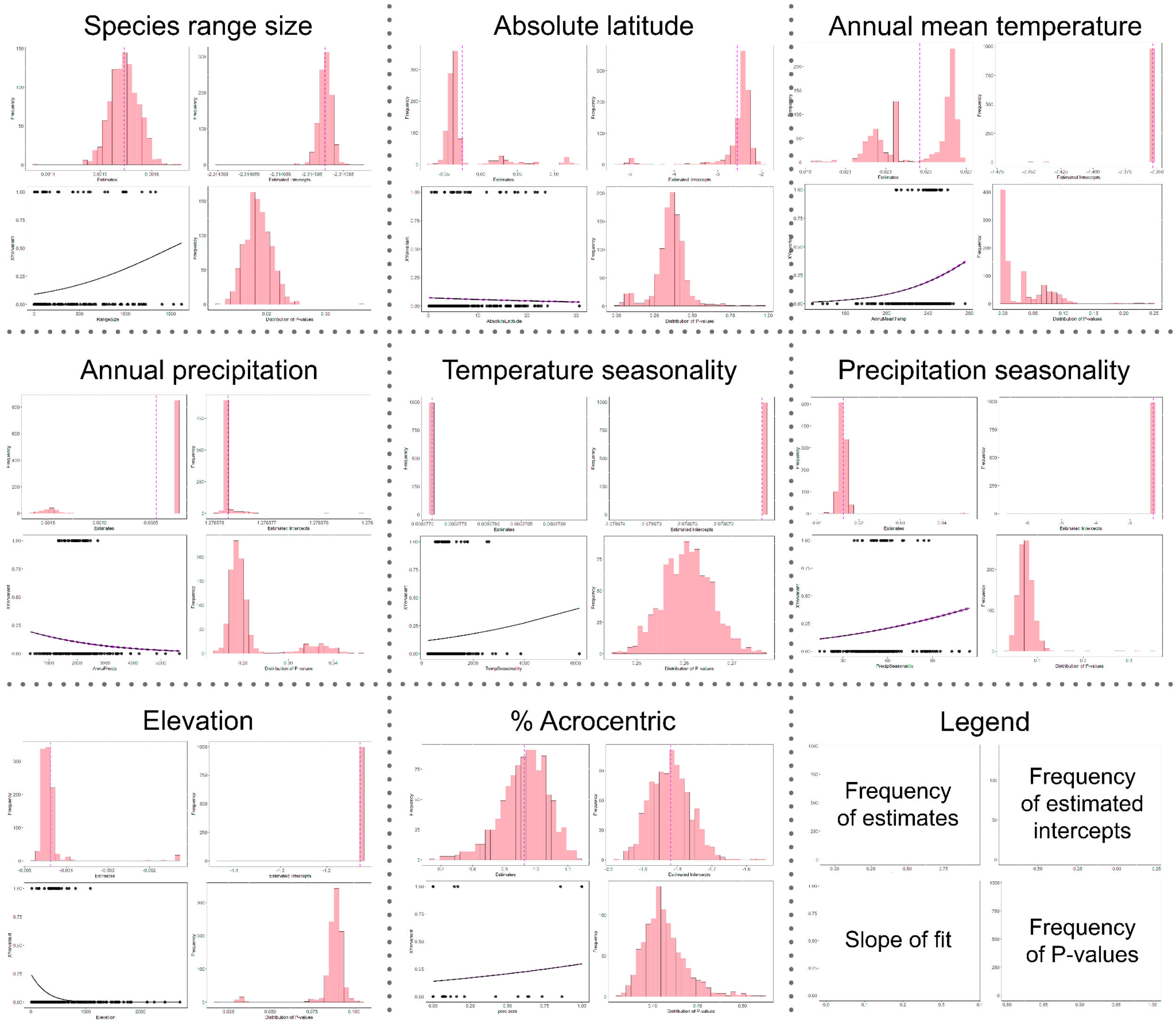


Supplemental Figure 1-3: *sensiPhy* hypothesis testing results of all eight predictor variables analyzed via phylogenetic logistic regression for all three mammal families of interest in this study, produced using the package's sensi_plot function. Vertical dashed lines in individual histograms indicate the mean estimate. Legend on lower right provides guidance on how to interpret histograms. Note the variability in the distribution of many of these summary statistics.

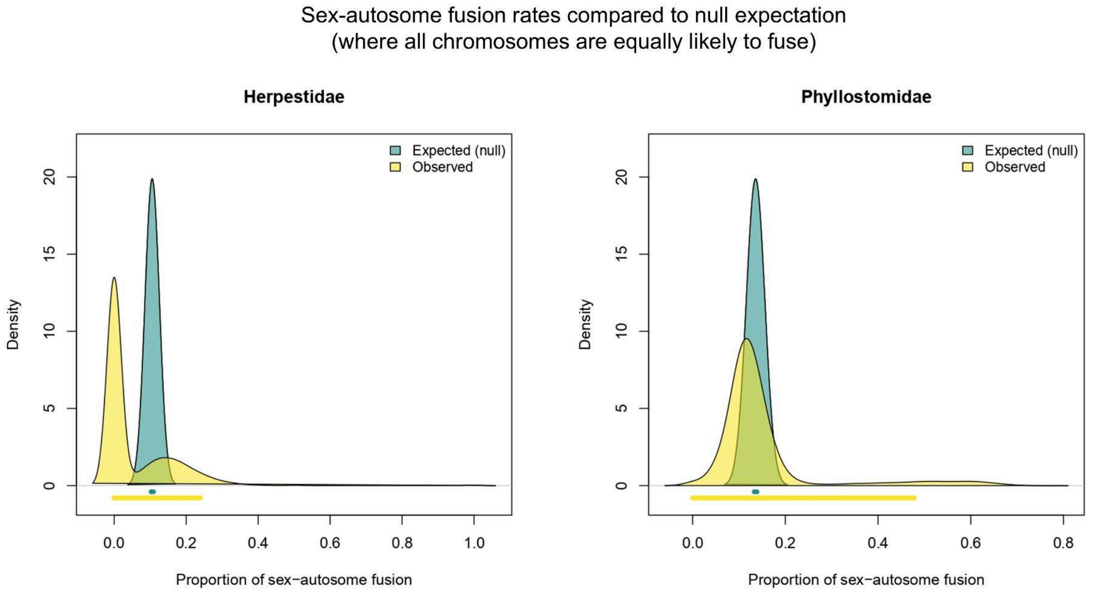


Supplemental Figure 4: Comparison of sex-autosome fusion rates against null expectations, where every chromosome is equally likely to undergo fusion. The 95% credible interval is shown by the line beneath the probability distribution. Each shows the rates as inferred across 1000 credible trees. Soricidae was not included in this analysis due to the single putative sex-autosome fusion event.

a)

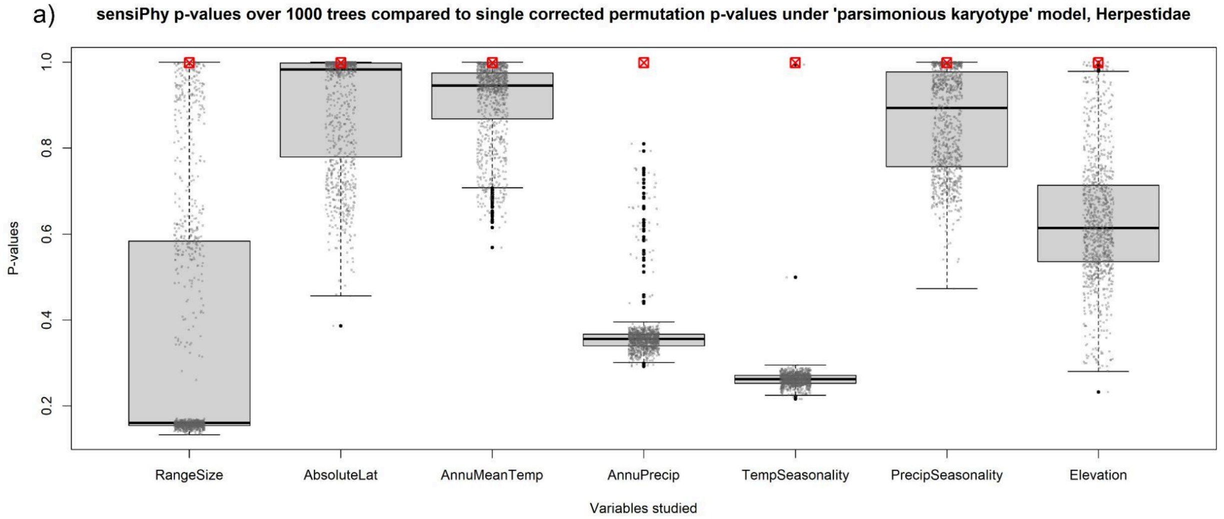


b)

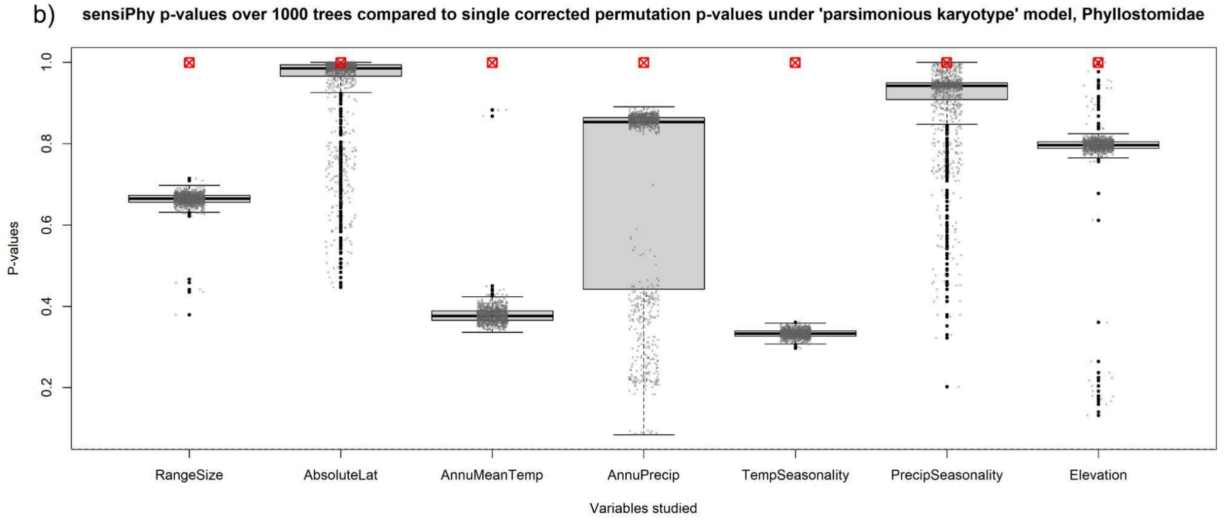


Supplemental Figure 5: Box-and-whisker plots of p-values from 1000 hypothesis tests conducted over our variables of interest using *sensiPhy* in our "Parsimonious Karyotype" dataset for a) Herpestidae and b) Phyllostomidae. Overlaid gray points are scatterplots of raw p-values from the same tests. Solid black points represent outlier data points in the box-and-whisker plots. Boxed red X's indicate permutation p-values for each test calculated via the formula $1 - [(\Sigma(number\ of\ tests\ with\ pval \leq 0.05) + 1) \div (1000 + 1)]$ after false discovery rate correction. Gray dashed line indicates the p≤0.05 threshold.

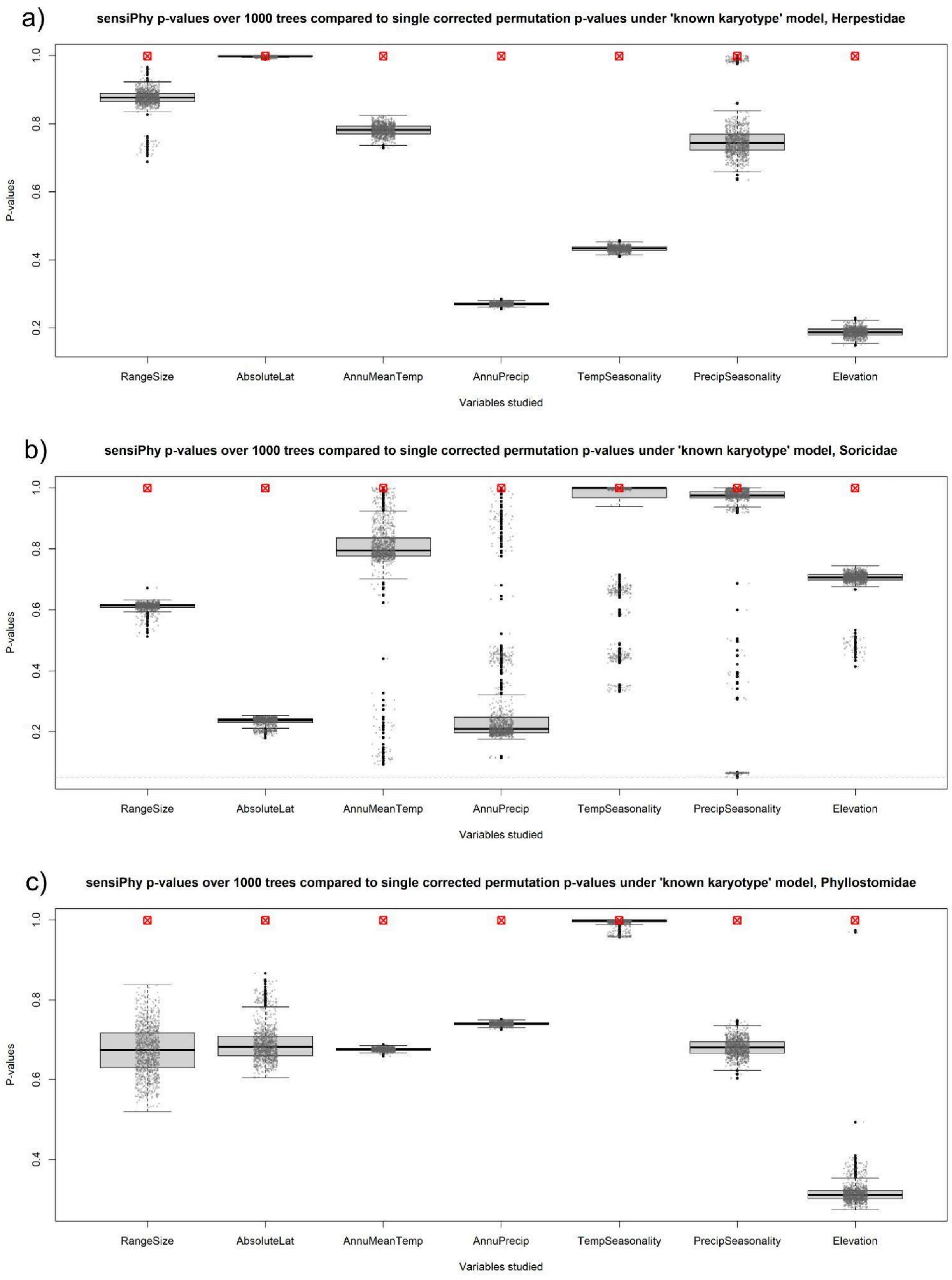
a)
sensiPhy p-values over 1000 trees compared to single corrected permutation p-values under 'known karyotype' model, Herpestidae
P-values
1.0
0.8
0.6
0.4
0.2
RangeSize
AbsoluteLat
AnnuMeanTemp
AnnuPrecip
TempSeasonality
PrecipSeasonality
Elevation
Variables studied
b)
sensiPhy p-values over 1000 trees compared to single corrected permutation p-values under 'known karyotype' model, Soricidae
P-values
1.0
0.8
0.6
0.4
0.2
RangeSize
AbsoluteLat
AnnuMeanTemp
AnnuPrecip
TempSeasonality
PrecipSeasonality
Elevation
Variables studied
c)
sensiPhy p-values over 1000 trees compared to single corrected permutation p-values under 'known karyotype' model, Phyllostomidae
P-values
1.0
0.8
0.6
0.4
RangeSize
AbsoluteLat
AnnuMeanTemp
AnnuPrecip
TempSeasonality
PrecipSeasonality
Elevation
Variables studied

Supplemental Figure 6: Box-and-whisker plots of p-values from 1000 hypothesis tests conducted over our variables of interest using *sensiPhy* in our “Known Karyotype” dataset for a) Herpestidae, b) Soricidae, and c) Phyllostomidae. Overlaid gray points are scatterplots of raw p-values from the same tests. Solid black points represent outlier data points in the box-and-whisker plots. Boxed red X’s indicate permutation p-values for each test calculated via the formula $1 - [(\Sigma(number\ of\ tests\ with\ pval \leq 0.05) + 1) \div (1000 + 1)]$ after false discovery rate correction. Gray dashed line indicates the p≤0.05 threshold.
