## Supplementary Table for "Ecoclimatic variables and karyotype features are poor predictors of neo-sex chromosome evolution in mammals"

**Supplemental Materials**

Supplemental Table 1: Comprehensive list of mammal species included in our analysis, sampled from three mammalian families. Of 34 total species from Herpestidae included in our analysis (excluding *Urva auropunctata*), 9 species were coded as possessing a Y-autosome fusion. Of 413 total species from Soricidae (excluding *Sorex kozlovi*), 10 species were coded as possessing an X-autosome fusion. Of 205 total species from Phyllostomidae, 15 species were coded as possessing an X-autosome fusion and 14 species were coded as possessing both an X- and Y-autosome fusion, summing to 29 variant species total.

| Family | Species | Sex | Known or assumed karyotype? | Citation | Comments |
|---|---|---|---|---|---|
| Herpestidae | *Atilax paludinosus* | YAfusion | Known | Blackmon et al. (2019) | NA |
| Herpestidae | *Bdeogale crassicauda* | XY | Assumed | NA | NA |
| Herpestidae | *Bdeogale jacksoni* | XY | Assumed | NA | NA |
| Herpestidae | *Bdeogale nigripes* | XY | Known | Wurster and Benirschke (1968) | NA |
| Herpestidae | *Bdeogale omnivora* | XY | Assumed | NA | NA |
| Herpestidae | *Crossarchus alexandri* | XY | Known | Perez et al. (2006) | While this karyotype has been studied, certain chromosome morphology attributes relevant to our study were not recorded. |
| Herpestidae | *Crossarchus ansorgei* | XY | Assumed | NA | NA |
| Herpestidae | *Crossarchus obscurus* | XY | Known | Blackmon et al. (2019) | NA |
| Herpestidae | *Crossarchus platycephalus* | XY | Assumed | NA | NA |
| Herpestidae | *Cynictis penicillata* | XY | Known | Blackmon et al. (2019) | NA |

| | | | | | |
|---|---|---|---|---|---|
| Herpestidae | *Dologale dybowskii* | XY | Assumed | NA | NA |
| Herpestidae | *Helogale hirtula* | XY | Known | Perez et al. (2006) | While this karyotype has been studied, certain chromosome morphology attributes relevant to our study were not recorded. |
| Herpestidae | *Helogale parvula* | XY | Known | Blackmon et al. (2019) | NA |
| Herpestidae | *Herpestes flavescens* | XY | Assumed | NA | Considered a variant sex chromosome species in “Parsimonious Karyotype” dataset. |
| Herpestidae | *Herpestes ichneumon* | YAfusion | Known | Blackmon et al. (2019) | NA |
| Herpestidae | *Herpestes ochraceus* | XY | Assumed | NA | Considered a variant sex chromosome species in “Parsimonious Karyotype” dataset. |
| Herpestidae | *Herpestes pulverulentus* | YAfusion | Known | Blackmon et al. (2019) | NA |
| Herpestidae | *Herpestes sanguineus* | YAfusion | Known | Fredga (1972) | NA |
| Herpestidae | *Ichneumia albicauda* | XY | Known | Wurster and Benirschke (1968) | NA |
| Herpestidae | *Liberiictis kuhni* | XY | Assumed | NA | NA |
| Herpestidae | *Mungos gambianus* | XY | Assumed | NA | NA |
| Herpestidae | *Mungos mungo* | XY | Known | Blackmon et al. (2019) | NA |
| Herpestidae | *Paracynictis selousi* | XY | Assumed | NA | NA |
| Herpestidae | *Rhynchogale melleri* | XY | Known | Perez et al. (2006) | While this karyotype has been studied, certain chromosome |

| | | | | | |
|---|---|---|---|---|---|
| | | | | | morphology attributes relevant to our study were not recorded. |
| Herpestidae | *Suricata suricatta* | XY | Known | Wurster and Benirschke (1968) | NA |
| Herpestidae | *Urva auropunctata* | YAfusion | Known | Fredga (1972), Murata et al. (2016) | Due to the lack of Urva auropunctata (small Indian mongoose) in the mammal phylogeny from Upham et al. (2019), we considered it synonymous with Urva javanica (Javan mongoose) given their putatively identical karyotypes, including a shared Y-autosome fusion, according to Fredga (1972). |
| Herpestidae | *Urva brachyura* | YAfusion | Known | Fredga (1972) | NA |
| Herpestidae | *Urva edwardsii* | YAfusion | Known | Fredga (1972) | NA |
| Herpestidae | *Urva fusca* | YAfusion | Known | Fredga (1972) | NA |
| Herpestidae | *Urva javanica* | YAfusion | Known | Blackmon et al. (2019) | NA |
| Herpestidae | *Urva semitorquata* | XY | Assumed | NA | Considered a variant sex chromosome species in “Parsimonious Karyotype” dataset. |
| Herpestidae | *Urva smithii* | XY | Assumed | NA | NA |
| Herpestidae | *Urva urva* | YAfusion | Known | Fredga (1972) | NA |
| Herpestidae | *Urva vitticollis* | XY | Assumed | NA | Considered a variant sex chromosome species in “Parsimonious |

| | | | | | |
|---|---|---|---|---|---|
| | | | | | Karyotype" dataset. |
| Herpestidae | *Xenogale naso* | XY | Assumed | NA | Considered a variant sex chromosome species in "Parsimonious Karyotype" dataset. |
| Soricidae | *Anourosorex assamensis* | XY | Assumed | NA | NA |
| Soricidae | *Anourosorex schmidi* | XY | Assumed | NA | NA |
| Soricidae | *Anourosorex squamipes* | XY | Known | Blackmon et al. (2019) | NA |
| Soricidae | *Anourosorex yamashinai* | XY | Assumed | NA | NA |
| Soricidae | *Blarina brevicauda* | XY | Known | Blackmon et al. (2019) | NA |
| Soricidae | *Blarina carolinensis* | XY | Known | Blackmon et al. (2019) | NA |
| Soricidae | *Blarina hylophaga* | XY | Known | Blackmon et al. (2019) | NA |
| Soricidae | *Blarinella griselda* | XY | Assumed | NA | NA |
| Soricidae | *Blarinella quadraticauda* | XY | Assumed | NA | NA |
| Soricidae | *Blarinella wardi* | XY | Assumed | NA | NA |
| Soricidae | *Chimarrogale hantu* | XY | Assumed | NA | NA |
| Soricidae | *Chimarrogale himalayica* | XY | Assumed | NA | NA |
| Soricidae | *Chimarrogale phaeura* | XY | Assumed | NA | NA |
| Soricidae | *Chimarrogale platycephalus* | XY | Assumed | NA | NA |
| Soricidae | *Chimarrogale styani* | XY | Assumed | NA | NA |
| Soricidae | *Chimarrogale sumatrana* | XY | Assumed | NA | NA |
| Soricidae | *Chodsigoa caovansunga* | XY | Assumed | NA | NA |
| Soricidae | *Chodsigoa hypsibia* | XY | Assumed | NA | NA |
| Soricidae | *Chodsigoa lamula* | XY | Assumed | NA | NA |

| | | | | | |
|---|---|---|---|---|---|
| Soricidae | *Chodsigoa parca* | XY | Assumed | NA | NA |
| Soricidae | *Chodsigoa parva* | XY | Assumed | NA | NA |
| Soricidae | *Chodsigoa salenskii* | XY | Assumed | NA | NA |
| Soricidae | *Chodsigoa smithii* | XY | Assumed | NA | NA |
| Soricidae | *Chodsigoa sodalis* | XY | Assumed | NA | NA |
| Soricidae | *Congosorex phillipsorum* | XY | Assumed | NA | NA |
| Soricidae | *Congosorex polli* | XY | Assumed | NA | NA |
| Soricidae | *Congosorex verheyeni* | XY | Assumed | NA | NA |
| Soricidae | *Crocidura abscondita* | XY | Assumed | NA | NA |
| Soricidae | *Crocidura aleksandrisi* | XY | Assumed | NA | NA |
| Soricidae | *Crocidura allex* | XY | Assumed | NA | NA |
| Soricidae | *Crocidura andamanensis* | XY | Assumed | NA | NA |
| Soricidae | *Crocidura annamitensis* | XY | Assumed | NA | NA |
| Soricidae | *Crocidura ansellorum* | XY | Assumed | NA | NA |
| Soricidae | *Crocidura arabica* | XY | Assumed | NA | NA |
| Soricidae | *Crocidura arispa* | XY | Assumed | NA | NA |
| Soricidae | *Crocidura armenica* | XY | Assumed | NA | NA |
| Soricidae | *Crocidura attenuata* | XY | Assumed | NA | NA |
| Soricidae | *Crocidura attila* | XY | Assumed | NA | NA |
| Soricidae | *Crocidura baileyi* | XY | Assumed | NA | NA |
| Soricidae | *Crocidura baluensis* | XY | Assumed | NA | NA |
| Soricidae | *Crocidura batakorum* | XY | Assumed | NA | NA |
| Soricidae | *Crocidura batesi* | XY | Assumed | NA | NA |

| | | | | | |
|---|---|---|---|---|---|
| Soricidae | *Crocidura beatus* | XY | Assumed | NA | NA |
| Soricidae | *Crocidura beccarii* | XY | Assumed | NA | NA |
| Soricidae | *Crocidura bottegi* | XY | Known | Blackmon et al. (2019) | NA |
| Soricidae | *Crocidura bottegoides* | XY | Assumed | NA | NA |
| Soricidae | *Crocidura brunnea* | XY | Assumed | NA | NA |
| Soricidae | *Crocidura buettikoferi* | XY | Assumed | NA | NA |
| Soricidae | *Crocidura caliginea* | XY | Assumed | NA | NA |
| Soricidae | *Crocidura canariensis* | XY | Assumed | NA | NA |
| Soricidae | *Crocidura caspica* | XY | Assumed | NA | NA |
| Soricidae | *Crocidura cinderella* | XY | Assumed | NA | NA |
| Soricidae | *Crocidura congobelgica* | XY | Assumed | NA | NA |
| Soricidae | *Crocidura cranbrooki* | XY | Assumed | NA | NA |
| Soricidae | *Crocidura crenata* | XY | Assumed | NA | NA |
| Soricidae | *Crocidura crossei* | XY | Known | Blackmon et al. (2019) | NA |
| Soricidae | *Crocidura cyanea* | XY | Assumed | NA | NA |
| Soricidae | *Crocidura denti* | XY | Assumed | NA | NA |
| Soricidae | *Crocidura desperata* | XY | Assumed | NA | NA |
| Soricidae | *Crocidura dhofarensis* | XY | Assumed | NA | NA |
| Soricidae | *Crocidura dolichura* | XY | Assumed | NA | NA |
| Soricidae | *Crocidura douceti* | XY | Assumed | NA | NA |
| Soricidae | *Crocidura dsinezumi* | XY | Assumed | NA | NA |
| Soricidae | *Crocidura eisentrauti* | XY | Assumed | NA | NA |
| Soricidae | *Crocidura elgonius* | XY | Assumed | NA | NA |

| | | | | | |
|---|---|---|---|---|---|
| Soricidae | *Crocidura elongata* | XY | Assumed | NA | NA |
| Soricidae | *Crocidura erica* | XY | Assumed | NA | NA |
| Soricidae | *Crocidura fingui* | XY | Assumed | NA | NA |
| Soricidae | *Crocidura fischeri* | XY | Assumed | NA | NA |
| Soricidae | *Crocidura flavescens* | XY | Assumed | NA | NA |
| Soricidae | *Crocidura floweri* | XY | Assumed | NA | NA |
| Soricidae | *Crocidura foetida* | XY | Assumed | NA | NA |
| Soricidae | *Crocidura foxi* | XY | Assumed | NA | NA |
| Soricidae | *Crocidura fuliginosa* | XY | Assumed | NA | NA |
| Soricidae | *Crocidura fulvastra* | XY | Assumed | NA | NA |
| Soricidae | *Crocidura fumosa* | XY | Assumed | NA | NA |
| Soricidae | *Crocidura fuscomurina* | XY | Assumed | NA | NA |
| Soricidae | *Crocidura glassi* | XY | Assumed | NA | NA |
| Soricidae | *Crocidura gmelini* | XY | Assumed | NA | NA |
| Soricidae | *Crocidura goliath* | XY | Assumed | NA | NA |
| Soricidae | *Crocidura gracilipes* | XY | Known | Blackmon et al. (2019) | NA |
| Soricidae | *Crocidura grandiceps* | XY | Assumed | NA | NA |
| Soricidae | *Crocidura grandis* | XY | Assumed | NA | NA |
| Soricidae | *Crocidura grassei* | XY | Assumed | NA | NA |
| Soricidae | *Crocidura grayi* | XY | Assumed | NA | NA |
| Soricidae | *Crocidura greenwoodi* | XY | Assumed | NA | NA |
| Soricidae | *Crocidura guy* | XY | Assumed | NA | NA |
| Soricidae | *Crocidura harenna* | XY | Assumed | NA | NA |
| Soricidae | *Crocidura hikmiya* | XY | Assumed | NA | NA |

| | | | | | |
|---|---|---|---|---|---|
| Soricidae | *Crocidura hildegardeae* | XY | Assumed | NA | NA |
| Soricidae | *Crocidura hilliana* | XY | Assumed | NA | NA |
| Soricidae | *Crocidura hirta* | XY | Assumed | NA | NA |
| Soricidae | *Crocidura hispida* | XY | Assumed | NA | NA |
| Soricidae | *Crocidura horsfieldii* | XY | Assumed | NA | NA |
| Soricidae | *Crocidura hutanis* | XY | Assumed | NA | NA |
| Soricidae | *Crocidura indochinensis* | XY | Assumed | NA | NA |
| Soricidae | *Crocidura jacksoni* | XY | Assumed | NA | NA |
| Soricidae | *Crocidura jenkinsi* | XY | Assumed | NA | NA |
| Soricidae | *Crocidura jouvenetae* | XY | Assumed | NA | NA |
| Soricidae | *Crocidura katinka* | XY | Assumed | NA | NA |
| Soricidae | *Crocidura kivuana* | XY | Assumed | NA | NA |
| Soricidae | *Crocidura lamottei* | XY | Known | Blackmon et al. (2019) | NA |
| Soricidae | *Crocidura lanosa* | XY | Assumed | NA | NA |
| Soricidae | *Crocidura lasiura* | XY | Known | Blackmon et al. (2019) | NA |
| Soricidae | *Crocidura latona* | XY | Assumed | NA | NA |
| Soricidae | *Crocidura lea* | XY | Assumed | NA | NA |
| Soricidae | *Crocidura lepidura* | XY | Assumed | NA | NA |
| Soricidae | *Crocidura leucodon* | XY | Known | Blackmon et al. (2019) | NA |
| Soricidae | *Crocidura levicula* | XY | Assumed | NA | NA |
| Soricidae | *Crocidura littoralis* | XY | Assumed | NA | NA |
| Soricidae | *Crocidura longipes* | XY | Assumed | NA | NA |
| Soricidae | *Crocidura lucina* | XY | Assumed | NA | NA |
| Soricidae | *Crocidura ludia* | XY | Assumed | NA | NA |

| | | | | | |
|---|---|---|---|---|---|
| Soricidae | *Crocidura luna* | XY | Assumed | NA | NA |
| Soricidae | *Crocidura lusitania* | XY | Assumed | NA | NA |
| Soricidae | *Crocidura lwiroensis* | XY | Assumed | NA | NA |
| Soricidae | *Crocidura macarthuri* | XY | Assumed | NA | NA |
| Soricidae | *Crocidura macmillani* | XY | Assumed | NA | NA |
| Soricidae | *Crocidura macowi* | XY | Assumed | NA | NA |
| Soricidae | *Crocidura malayana* | XY | Assumed | NA | NA |
| Soricidae | *Crocidura manengubae* | XY | Assumed | NA | NA |
| Soricidae | *Crocidura maquassiensis* | XY | Assumed | NA | NA |
| Soricidae | *Crocidura mariquensis* | XY | Assumed | NA | NA |
| Soricidae | *Crocidura maurisca* | XY | Assumed | NA | NA |
| Soricidae | *Crocidura maxi* | XY | Assumed | NA | NA |
| Soricidae | *Crocidura mdumai* | XY | Assumed | NA | NA |
| Soricidae | *Crocidura mindorus* | XY | Assumed | NA | NA |
| Soricidae | *Crocidura miya* | XY | Assumed | NA | NA |
| Soricidae | *Crocidura monax* | XY | Assumed | NA | NA |
| Soricidae | *Crocidura monticola* | XY | Assumed | NA | NA |
| Soricidae | *Crocidura montis* | XY | Assumed | NA | NA |
| Soricidae | *Crocidura munissii* | XY | Assumed | NA | NA |
| Soricidae | *Crocidura muricauda* | XY | Assumed | NA | NA |
| Soricidae | *Crocidura musseri* | XY | Assumed | NA | NA |
| Soricidae | *Crocidura mutesae* | XY | Assumed | NA | NA |
| Soricidae | *Crocidura nana* | XY | Assumed | NA | NA |
| Soricidae | *Crocidura nanilla* | XY | Assumed | NA | NA |

| | | | | | |
|---|---|---|---|---|---|
| Soricidae | *Crocidura neglecta* | XY | Assumed | NA | NA |
| Soricidae | *Crocidura negligens* | XY | Assumed | NA | NA |
| Soricidae | *Crocidura negrina* | XY | Assumed | NA | NA |
| Soricidae | *Crocidura newmarki* | XY | Assumed | NA | NA |
| Soricidae | *Crocidura nicobarica* | XY | Assumed | NA | NA |
| Soricidae | *Crocidura nigeriae* | XY | Known | Blackmon et al. (2019) | NA |
| Soricidae | *Crocidura nigricans* | XY | Assumed | NA | NA |
| Soricidae | *Crocidura nigripes* | XY | Assumed | NA | NA |
| Soricidae | *Crocidura nigrofusca* | XY | Assumed | NA | NA |
| Soricidae | *Crocidura nimbae* | XY | Known | Blackmon et al. (2019) | NA |
| Soricidae | *Crocidura nimbasilvanus* | XY | Assumed | NA | NA |
| Soricidae | *Crocidura ninoyi* | XY | Assumed | NA | NA |
| Soricidae | *Crocidura niobe* | XY | Assumed | NA | NA |
| Soricidae | *Crocidura obscurior* | XY | Assumed | NA | NA |
| Soricidae | *Crocidura olivieri* | XY | Assumed | NA | NA |
| Soricidae | *Crocidura orientalis* | XY | Assumed | NA | NA |
| Soricidae | *Crocidura orii* | XY | Assumed | NA | NA |
| Soricidae | *Crocidura pachyura* | XY | Assumed | NA | NA |
| Soricidae | *Crocidura palawanensis* | XY | Assumed | NA | NA |
| Soricidae | *Crocidura panayensis* | XY | Assumed | NA | NA |
| Soricidae | *Crocidura paradoxura* | XY | Assumed | NA | NA |
| Soricidae | *Crocidura parvipes* | XY | Assumed | NA | NA |
| Soricidae | *Crocidura pasha* | XY | Assumed | NA | NA |

| | | | | | |
|---|---|---|---|---|---|
| Soricidae | *Crocidura pergrisea* | XY | Assumed | NA | NA |
| Soricidae | *Crocidura phaeura* | XY | Assumed | NA | NA |
| Soricidae | *Crocidura phanluongi* | XY | Assumed | NA | NA |
| Soricidae | *Crocidura phuquocensis* | XY | Assumed | NA | NA |
| Soricidae | *Crocidura picea* | XY | Assumed | NA | NA |
| Soricidae | *Crocidura pitmani* | XY | Assumed | NA | NA |
| Soricidae | *Crocidura planiceps* | XY | Known | Blackmon et al. (2019) | NA |
| Soricidae | *Crocidura poensis* | XY | Known | Blackmon et al. (2019) | NA |
| Soricidae | *Crocidura polia* | XY | Assumed | NA | NA |
| Soricidae | *Crocidura pullata* | XY | Assumed | NA | NA |
| Soricidae | *Crocidura raineyi* | XY | Assumed | NA | NA |
| Soricidae | *Crocidura ramona* | XY | Assumed | NA | NA |
| Soricidae | *Crocidura rapax* | XY | Assumed | NA | NA |
| Soricidae | *Crocidura religiosa* | XY | Assumed | NA | NA |
| Soricidae | *Crocidura rhoditis* | XY | Assumed | NA | NA |
| Soricidae | *Crocidura roosevelti* | XY | Assumed | NA | NA |
| Soricidae | *Crocidura russula* | XY | Known | Blackmon et al. (2019) | NA |
| Soricidae | *Crocidura sapaensis* | XY | Assumed | NA | NA |
| Soricidae | *Crocidura selina* | XY | Assumed | NA | NA |
| Soricidae | *Crocidura serezkyensis* | XY | Assumed | NA | NA |
| Soricidae | *Crocidura shantungensis* | XY | Assumed | NA | NA |
| Soricidae | *Crocidura sibirica* | XY | Assumed | NA | NA |
| Soricidae | *Crocidura sicula* | XY | Assumed | NA | NA |

| | | | | | |
|---|---|---|---|---|---|
| Soricidae | *Crocidura silacea* | XY | Assumed | NA | NA |
| Soricidae | *Crocidura smithii* | XY | Assumed | NA | NA |
| Soricidae | *Crocidura sokolovi* | XY | Assumed | NA | NA |
| Soricidae | *Crocidura somalica* | XY | Assumed | NA | NA |
| Soricidae | *Crocidura stenocephala* | XY | Assumed | NA | NA |
| Soricidae | *Crocidura suaveolens* | XY | Known | Blackmon et al. (2019) | NA |
| Soricidae | *Crocidura susiana* | XY | Assumed | NA | NA |
| Soricidae | *Crocidura tanakae* | XY | Assumed | NA | NA |
| Soricidae | *Crocidura tansaniana* | XY | Assumed | NA | NA |
| Soricidae | *Crocidura tarella* | XY | Assumed | NA | NA |
| Soricidae | *Crocidura tarfayensis* | XY | Assumed | NA | NA |
| Soricidae | *Crocidura telfordi* | XY | Assumed | NA | NA |
| Soricidae | *Crocidura tenuis* | XY | Assumed | NA | NA |
| Soricidae | *Crocidura thalia* | XY | Assumed | NA | NA |
| Soricidae | *Crocidura theresae* | XY | Known | Blackmon et al. (2019) | NA |
| Soricidae | *Crocidura thomensis* | XY | Assumed | NA | NA |
| Soricidae | *Crocidura trichura* | XY | Assumed | NA | NA |
| Soricidae | *Crocidura turba* | XY | Assumed | NA | NA |
| Soricidae | *Crocidura ultima* | XY | Assumed | NA | NA |
| Soricidae | *Crocidura usambarae* | XY | Assumed | NA | NA |
| Soricidae | *Crocidura viaria* | XY | Assumed | NA | NA |
| Soricidae | *Crocidura virgata* | XY | Assumed | NA | NA |
| Soricidae | *Crocidura voi* | XY | Assumed | NA | NA |

| | | | | | |
|---|---|---|---|---|---|
| Soricidae | *Crocidura vorax* | XY | Assumed | NA | NA |
| Soricidae | *Crocidura vosmaeri* | XY | Assumed | NA | NA |
| Soricidae | *Crocidura watasei* | XY | Assumed | NA | NA |
| Soricidae | *Crocidura whitakeri* | XY | Assumed | NA | NA |
| Soricidae | *Crocidura wimmeri* | XY | Known | Blackmon et al. (2019) | NA |
| Soricidae | *Crocidura wuchihensis* | XY | Assumed | NA | NA |
| Soricidae | *Crocidura xantippe* | XY | Assumed | NA | NA |
| Soricidae | *Crocidura yankariensis* | XY | Assumed | NA | NA |
| Soricidae | *Crocidura zaitsevi* | XY | Assumed | NA | NA |
| Soricidae | *Crocidura zaphiri* | XY | Assumed | NA | NA |
| Soricidae | *Crocidura zarudnyi* | XY | Assumed | NA | NA |
| Soricidae | *Crocidura zimmeri* | XY | Assumed | NA | NA |
| Soricidae | *Crocidura zimmermanni* | XY | Known | Blackmon et al. (2019) | NA |
| Soricidae | *Cryptotis alticola* | XY | Assumed | NA | NA |
| Soricidae | *Cryptotis aroensis* | XY | Assumed | NA | NA |
| Soricidae | *Cryptotis brachyonyx* | XY | Assumed | NA | NA |
| Soricidae | *Cryptotis colombiana* | XY | Assumed | NA | NA |
| Soricidae | *Cryptotis endersi* | XY | Assumed | NA | NA |
| Soricidae | *Cryptotis equatoris* | XY | Assumed | NA | NA |
| Soricidae | *Cryptotis goldmani* | XY | Assumed | NA | NA |
| Soricidae | *Cryptotis goodwini* | XY | Assumed | NA | NA |
| Soricidae | *Cryptotis gracilis* | XY | Assumed | NA | NA |

| | | | | | |
|---|---|---|---|---|---|
| Soricidae | *Cryptotis griseoventris* | XY | Assumed | NA | NA |
| Soricidae | *Cryptotis hondurensis* | XY | Assumed | NA | NA |
| Soricidae | *Cryptotis lacandonensis* | XY | Assumed | NA | NA |
| Soricidae | *Cryptotis lacertosus* | XY | Assumed | NA | NA |
| Soricidae | *Cryptotis magna* | XY | Assumed | NA | NA |
| Soricidae | *Cryptotis mam* | XY | Assumed | NA | NA |
| Soricidae | *Cryptotis mayensis* | XY | Assumed | NA | NA |
| Soricidae | *Cryptotis medellinia* | XY | Assumed | NA | NA |
| Soricidae | *Cryptotis mera* | XY | Assumed | NA | NA |
| Soricidae | *Cryptotis meridensis* | XY | Assumed | NA | NA |
| Soricidae | *Cryptotis merriami* | XY | Assumed | NA | NA |
| Soricidae | *Cryptotis mexicana* | XY | Assumed | NA | NA |
| Soricidae | *Cryptotis montivaga* | XY | Assumed | NA | NA |
| Soricidae | *Cryptotis nelsoni* | XY | Assumed | NA | NA |
| Soricidae | *Cryptotis niausa* | XY | Assumed | NA | NA |
| Soricidae | *Cryptotis nigrescens* | XY | Assumed | NA | NA |
| Soricidae | *Cryptotis obscura* | XY | Assumed | NA | NA |
| Soricidae | *Cryptotis oreoryctes* | XY | Assumed | NA | NA |
| Soricidae | *Cryptotis orophila* | XY | Assumed | NA | NA |
| Soricidae | *Cryptotis parva* | XY | Known | Blackmon et al. (2019) | NA |
| Soricidae | *Cryptotis peregrina* | XY | Assumed | NA | NA |
| Soricidae | *Cryptotis perijensis* | XY | Assumed | NA | NA |
| Soricidae | *Cryptotis peruviensis* | XY | Assumed | NA | NA |

| | | | | | |
|---|---|---|---|---|---|
| Soricidae | *Cryptotis phillipsii* | XY | Assumed | NA | NA |
| Soricidae | *Cryptotis squamipes* | XY | Assumed | NA | NA |
| Soricidae | *Cryptotis tamensis* | XY | Assumed | NA | NA |
| Soricidae | *Cryptotis thomasi* | XY | Assumed | NA | NA |
| Soricidae | *Cryptotis tropicalis* | XY | Assumed | NA | NA |
| Soricidae | *Cryptotis venezuelensis* | XY | Assumed | NA | NA |
| Soricidae | *Diplomesodon pulchellum* | XY | Known | Blackmon et al. (2019) | NA |
| Soricidae | *Episoriculus caudatus* | XY | Assumed | NA | NA |
| Soricidae | *Episoriculus fumidus* | XY | Assumed | NA | NA |
| Soricidae | *Episoriculus leucops* | XY | Assumed | NA | NA |
| Soricidae | *Episoriculus macrurus* | XY | Assumed | NA | NA |
| Soricidae | *Feroculus feroculus* | XY | Assumed | NA | NA |
| Soricidae | *Megasorex gigas* | XY | Assumed | NA | NA |
| Soricidae | *Myosorex babaulti* | XY | Assumed | NA | NA |
| Soricidae | *Myosorex blarina* | XY | Assumed | NA | NA |
| Soricidae | *Myosorex bururiensis* | XY | Assumed | NA | NA |
| Soricidae | *Myosorex cafer* | XY | Assumed | NA | NA |
| Soricidae | *Myosorex eisentrauti* | XY | Assumed | NA | NA |
| Soricidae | *Myosorex geata* | XY | Assumed | NA | NA |
| Soricidae | *Myosorex gnoskei* | XY | Assumed | NA | NA |
| Soricidae | *Myosorex jejei* | XY | Assumed | NA | NA |
| Soricidae | *Myosorex kabogoensis* | XY | Assumed | NA | NA |
| Soricidae | *Myosorex kihaulei* | XY | Assumed | NA | NA |

| | | | | | |
|---|---|---|---|---|---|
| Soricidae | *Myosorex longicaudatus* | XY | Assumed | NA | NA |
| Soricidae | *Myosorex meesteri* | XY | Assumed | NA | NA |
| Soricidae | *Myosorex okuensis* | XY | Assumed | NA | NA |
| Soricidae | *Myosorex rumpii* | XY | Assumed | NA | NA |
| Soricidae | *Myosorex schalleri* | XY | Assumed | NA | NA |
| Soricidae | *Myosorex sclateri* | XY | Assumed | NA | NA |
| Soricidae | *Myosorex tenuis* | XY | Assumed | NA | NA |
| Soricidae | *Myosorex varius* | XY | Assumed | NA | NA |
| Soricidae | *Myosorex zinki* | XY | Assumed | NA | NA |
| Soricidae | *Nectogale elegans* | XY | Assumed | NA | NA |
| Soricidae | *Neomys anomalus* | XY | Known | Blackmon et al. (2019) | NA |
| Soricidae | *Neomys fodiens* | XY | Known | Blackmon et al. (2019) | NA |
| Soricidae | *Neomys teres* | XY | Assumed | NA | NA |
| Soricidae | *Notiosorex cockrumi* | XY | Assumed | NA | NA |
| Soricidae | *Notiosorex crawfordi* | XY | Assumed | NA | NA |
| Soricidae | *Notiosorex evotis* | XY | Assumed | NA | NA |
| Soricidae | *Notiosorex villai* | XY | Assumed | NA | NA |
| Soricidae | *Paracrocidura graueri* | XY | Assumed | NA | NA |
| Soricidae | *Paracrocidura maxima* | XY | Assumed | NA | NA |
| Soricidae | *Paracrocidura schoutedeni* | XY | Assumed | NA | NA |
| Soricidae | *Ruwenzorisorex suncoides* | XY | Assumed | NA | NA |
| Soricidae | *Scutisorex somereni* | XY | Assumed | NA | NA |
| Soricidae | *Scutisorex thori* | XY | Assumed | NA | NA |
| Soricidae | *Solisorex pearsoni* | XY | Assumed | NA | NA |

| | | | | | |
|---|---|---|---|---|---|
| Soricidae | *Sorex alaskanus* | XY | Assumed | NA | NA |
| Soricidae | *Sorex alpinus* | XY | Assumed | NA | NA |
| Soricidae | *Sorex antinorii* | XAfusion | Known | Bulatova et al. (2019) | NA |
| Soricidae | *Sorex araneus* | XAfusion | Known | Blackmon et al. (2019) | NA |
| Soricidae | *Sorex arcticus* | XAfusion | Known | Bulatova et al. (2019) | While this karyotype has been studied, certain chromosome morphology attributes relevant to our study were not recorded. |
| Soricidae | *Sorex arizonae* | XY | Assumed | NA | NA |
| Soricidae | *Sorex arunchi* | XY | Assumed | NA | NA |
| Soricidae | *Sorex asper* | XAfusion | Known | Blackmon et al. (2019) | NA |
| Soricidae | *Sorex bairdi* | XY | Assumed | NA | NA |
| Soricidae | *Sorex bedfordiae* | XY | Assumed | NA | NA |
| Soricidae | *Sorex bendirii* | XY | Known | Blackmon et al. (2019) | NA |
| Soricidae | *Sorex buchariensis* | XY | Known | Blackmon et al. (2019) | NA |
| Soricidae | *Sorex caecutiens* | XY | Known | Blackmon et al. (2019) | NA |
| Soricidae | *Sorex camtschatica* | XY | Assumed | NA | NA |
| Soricidae | *Sorex cansulus* | XY | Assumed | NA | NA |
| Soricidae | *Sorex cinereus* | XY | Known | Blackmon et al. (2019) | NA |
| Soricidae | *Sorex coronatus* | XAfusion | Known | Blackmon et al. (2019) | NA |
| Soricidae | *Sorex cylindricauda* | XY | Assumed | NA | NA |
| Soricidae | *Sorex daphaenodon* | XAfusion | Known | Blackmon et al. (2019) | NA |
| Soricidae | *Sorex dispar* | XY | Assumed | NA | NA |
| Soricidae | *Sorex emarginatus* | XY | Assumed | NA | NA |
| Soricidae | *Sorex excelsus* | XY | Assumed | NA | NA |

| | | | | | |
|---|---|---|---|---|---|
| Soricidae | *Sorex fumeus* | XY | Known | Blackmon et al. (2019) | NA |
| Soricidae | *Sorex gracillimus* | XY | Assumed | NA | NA |
| Soricidae | *Sorex granarius* | XAfusion | Known | Blackmon et al. (2019) | NA |
| Soricidae | *Sorex haydeni* | XY | Assumed | NA | NA |
| Soricidae | *Sorex hosonoi* | XY | Assumed | NA | NA |
| Soricidae | *Sorex hoyi* | XY | Assumed | NA | NA |
| Soricidae | *Sorex isodon* | XY | Known | Blackmon et al. (2019) | NA |
| Soricidae | *Sorex ixtlanensis* | XY | Assumed | NA | NA |
| Soricidae | *Sorex jacksoni* | XY | Assumed | NA | NA |
| Soricidae | *Sorex kozlovi* | XY | Assumed | NA | Sorex kozlovi is traditionally placed in the Sorex minutus group (Bannikova et al. 2018). In the Upham et al. (2019) phylogeny, its position is instead imputed as sister to Sorex daphaenodon as the only assumed XX/XY species in a clade otherwise wholly populated by species with X-autosome fusions. Given the lack of available cytological evidence to support or falsify this likely spurious placement, it was excluded from our trees and analyses. |
| Soricidae | *Sorex leucogaster* | XY | Assumed | NA | NA |

| | | | | | |
|---|---|---|---|---|---|
| Soricidae | *Sorex longirostris* | XY | Assumed | NA | NA |
| Soricidae | *Sorex lyelli* | XY | Assumed | NA | NA |
| Soricidae | *Sorex macrodon* | XY | Assumed | NA | NA |
| Soricidae | *Sorex maritimensis* | XAfusion | Known | Bulatova et al. (2019) | NA |
| Soricidae | *Sorex mediopua* | XY | Assumed | NA | NA |
| Soricidae | *Sorex merriami* | XY | Assumed | NA | NA |
| Soricidae | *Sorex milleri* | XY | Assumed | NA | NA |
| Soricidae | *Sorex minutissimus* | XY | Known | Blackmon et al. (2019) | NA |
| Soricidae | *Sorex minutus* | XY | Known | Blackmon et al. (2019) | NA |
| Soricidae | *Sorex mirabilis* | XY | Known | Blackmon et al. (2019) | NA |
| Soricidae | *Sorex monticolus* | XY | Assumed | NA | NA |
| Soricidae | *Sorex nanus* | XY | Assumed | NA | NA |
| Soricidae | *Sorex neomexicanus* | XY | Assumed | NA | NA |
| Soricidae | *Sorex oreopolus* | XY | Assumed | NA | NA |
| Soricidae | *Sorex orizabae* | XY | Assumed | NA | NA |
| Soricidae | *Sorex ornatus* | XY | Known | Blackmon et al. (2019) | NA |
| Soricidae | *Sorex pacificus* | XY | Assumed | NA | NA |
| Soricidae | *Sorex palustris* | XY | Assumed | NA | NA |
| Soricidae | *Sorex planiceps* | XY | Assumed | NA | NA |
| Soricidae | *Sorex portenkoi* | XY | Assumed | NA | NA |
| Soricidae | *Sorex preblei* | XY | Assumed | NA | NA |
| Soricidae | *Sorex pribilofensis* | XY | Assumed | NA | NA |
| Soricidae | *Sorex raddei* | XY | Known | Blackmon et al. (2019) | NA |
| Soricidae | *Sorex roboratus* | XY | Assumed | NA | NA |
| Soricidae | *Sorex rohweri* | XY | Assumed | NA | NA |
| Soricidae | *Sorex samniticus* | XY | Known | Blackmon et al. (2019) | NA |

| | | | | | |
|---|---|---|---|---|---|
| Soricidae | *Sorex satunini* | XAfusion | Known | Bulatova et al. (2019) | NA |
| Soricidae | *Sorex saussurei* | XY | Assumed | NA | NA |
| Soricidae | *Sorex sclateri* | XY | Assumed | NA | NA |
| Soricidae | *Sorex shinto* | XY | Known | Blackmon et al. (2019) | NA |
| Soricidae | *Sorex sinalis* | XY | Assumed | NA | NA |
| Soricidae | *Sorex sonomae* | XY | Assumed | NA | NA |
| Soricidae | *Sorex stizodon* | XY | Assumed | NA | NA |
| Soricidae | *Sorex tenellus* | XY | Assumed | NA | NA |
| Soricidae | *Sorex thibetanus* | XY | Assumed | NA | NA |
| Soricidae | *Sorex trowbridgii* | XY | Known | Blackmon et al. (2019) | NA |
| Soricidae | *Sorex tundrensis* | XAfusion | Known | Blackmon et al. (2019) | NA |
| Soricidae | *Sorex ugyunak* | XY | Assumed | NA | NA |
| Soricidae | *Sorex unguiculatus* | XY | Assumed | NA | NA |
| Soricidae | *Sorex vagrans* | XY | Known | Blackmon et al. (2019) | NA |
| Soricidae | *Sorex ventralis* | XY | Assumed | NA | NA |
| Soricidae | *Sorex veraecrucis* | XY | Assumed | NA | NA |
| Soricidae | *Sorex veraepacis* | XY | Assumed | NA | NA |
| Soricidae | *Sorex volnuchini* | XY | Assumed | NA | NA |
| Soricidae | *Sorex yukonicus* | XY | Assumed | NA | NA |
| Soricidae | *Soriculus nigrescens* | XY | Assumed | NA | NA |
| Soricidae | *Suncus aequatorius* | XY | Assumed | NA | NA |
| Soricidae | *Suncus ater* | XY | Assumed | NA | NA |
| Soricidae | *Suncus dayi* | XY | Assumed | NA | NA |
| Soricidae | *Suncus etruscus* | XY | Known | Blackmon et al. (2019) | NA |
| Soricidae | *Suncus fellowesgordoni* | XY | Assumed | NA | NA |
| Soricidae | *Suncus hosei* | XY | Assumed | NA | NA |
| Soricidae | *Suncus hututsi* | XY | Assumed | NA | NA |

| | | | | | |
|---|---|---|---|---|---|
| Soricidae | *Suncus infinitesimus* | XY | Assumed | NA | NA |
| Soricidae | *Suncus lixus* | XY | Assumed | NA | NA |
| Soricidae | *Suncus madagascariensis* | XY | Assumed | NA | NA |
| Soricidae | *Suncus malayanus* | XY | Assumed | NA | NA |
| Soricidae | *Suncus megalura* | XY | Assumed | NA | NA |
| Soricidae | *Suncus mertensi* | XY | Assumed | NA | NA |
| Soricidae | *Suncus montanus* | XY | Assumed | NA | NA |
| Soricidae | *Suncus murinus* | XY | Known | Blackmon et al. (2019) | NA |
| Soricidae | *Suncus remyi* | XY | Assumed | NA | NA |
| Soricidae | *Suncus stoliczkanus* | XY | Assumed | NA | NA |
| Soricidae | *Suncus varilla* | XY | Assumed | NA | NA |
| Soricidae | *Suncus zeylanicus* | XY | Assumed | NA | NA |
| Soricidae | *Surdisorex norae* | XY | Assumed | NA | NA |
| Soricidae | *Surdisorex polulus* | XY | Assumed | NA | NA |
| Soricidae | *Surdisorex schlitteri* | XY | Assumed | NA | NA |
| Soricidae | *Sylvisorex akaibei* | XY | Assumed | NA | NA |
| Soricidae | *Sylvisorex camerunensis* | XY | Assumed | NA | NA |
| Soricidae | *Sylvisorex corbeti* | XY | Assumed | NA | NA |
| Soricidae | *Sylvisorex granti* | XY | Assumed | NA | NA |
| Soricidae | *Sylvisorex howelli* | XY | Assumed | NA | NA |
| Soricidae | *Sylvisorex isabellae* | XY | Assumed | NA | NA |
| Soricidae | *Sylvisorex johnstoni* | XY | Assumed | NA | NA |
| Soricidae | *Sylvisorex konganensis* | XY | Assumed | NA | NA |

| | | | | | |
|---|---|---|---|---|---|
| Soricidae | *Sylvisorex lunaris* | XY | Assumed | NA | NA |
| Soricidae | *Sylvisorex morio* | XY | Assumed | NA | NA |
| Soricidae | *Sylvisorex ollula* | XY | Assumed | NA | NA |
| Soricidae | *Sylvisorex oriundus* | XY | Assumed | NA | NA |
| Soricidae | *Sylvisorex pluvialis* | XY | Assumed | NA | NA |
| Soricidae | *Sylvisorex silvanorum* | XY | Assumed | NA | NA |
| Soricidae | *Sylvisorex vulcanorum* | XY | Assumed | NA | NA |
| Phyllostomidae | *Ametrida centurio* | XAfusion | Known | Blackmon et al. (2019) | NA |
| Phyllostomidae | *Anoura canishina* | XY | Assumed | NA | NA |
| Phyllostomidae | *Anoura caudifer* | XY | Known | Baker et al. (1982) | NA |
| Phyllostomidae | *Anoura cultrata* | XY | Known | Baker et al. (1982) | NA |
| Phyllostomidae | *Anoura fistulata* | XY | Assumed | NA | NA |
| Phyllostomidae | *Anoura geoffroyi* | XY | Known | Baker et al. (1982) | NA |
| Phyllostomidae | *Anoura latidens* | XY | Assumed | NA | NA |
| Phyllostomidae | *Anoura luismanueli* | XY | Assumed | NA | NA |
| Phyllostomidae | *Ardops nichollsi* | XAfusion | Known | Blackmon et al. (2019) | NA |
| Phyllostomidae | *Ariteus flavescens* | XAfusion | Known | Greenbaum et al. (1975) | NA |
| Phyllostomidae | *Artibeus amplus* | XY | Assumed | NA | Considered a variant sex chromosome species in “Parsimonious Karyotype” dataset. |
| Phyllostomidae | *Artibeus concolor* | XY | Assumed | NA | Considered a variant sex chromosome species in “Parsimonious Karyotype” dataset. |

| | | | | | |
|---|---|---|---|---|---|
| Phyllostomidae | *Artibeus fimbriatus* | XAfusion | Known | Pinto et al. (2012) | NA |
| Phyllostomidae | *Artibeus fraterculus* | XY | Assumed | NA | Considered a variant sex chromosome species in “Parsimonious Karyotype” dataset. |
| Phyllostomidae | *Artibeus hirsutus* | XY | Assumed | NA | Considered a variant sex chromosome species in “Parsimonious Karyotype” dataset. |
| Phyllostomidae | *Artibeus inopinatus* | XY | Assumed | NA | Considered a variant sex chromosome species in “Parsimonious Karyotype” dataset. |
| Phyllostomidae | *Artibeus jamaicensis* | XAfusion | Known | Baker (1967) | NA |
| Phyllostomidae | *Artibeus lituratus* | XAfusion | Known | Yonenaga et al. (1969) | NA |
| Phyllostomidae | *Artibeus obscurus* | XAfusion | Known | Pieczarka et al. (2013) | NA |
| Phyllostomidae | *Artibeus planirostris* | XAfusion | Known | Pinto et al. (2012) | NA |
| Phyllostomidae | *Brachyphylla cavernarum* | XY | Assumed | NA | NA |
| Phyllostomidae | *Brachyphylla nana* | XY | Known | Blackmon et al. (2019) | NA |
| Phyllostomidae | *Carollia benkeithi* | XY | Assumed | NA | Considered a variant sex chromosome species in “Parsimonious Karyotype” dataset. |
| Phyllostomidae | *Carollia brevicauda* | XAfusion | Known | Blackmon et al. (2019) | NA |
| Phyllostomidae | *Carollia castanea* | XAfusion | Known | Blackmon et al. (2019) | NA |
| Phyllostomidae | *Carollia manu* | XY | Assumed | NA | Considered a variant sex chromosome |

| | | | | | |
|---|---|---|---|---|---|
| | | | | | species in “Parsimonious Karyotype” dataset. |
| Phyllostomidae | *Carollia perspicillata* | XAfusion | Known | Yonenaga et al. (1969) | NA |
| Phyllostomidae | *Carollia sowelli* | XY | Assumed | NA | Considered a variant sex chromosome species in “Parsimonious Karyotype” dataset. |
| Phyllostomidae | *Carollia subrufa* | XAfusion | Known | Blackmon et al. (2019) | NA |
| Phyllostomidae | *Centurio senex* | XY | Known | Blackmon et al. (2019) | NA |
| Phyllostomidae | *Chiroderma doriae* | XY | Assumed | NA | NA |
| Phyllostomidae | *Chiroderma improvisum* | XY | Known | Blackmon et al. (2019) | NA |
| Phyllostomidae | *Chiroderma salvini* | XY | Assumed | NA | NA |
| Phyllostomidae | *Chiroderma trinitatum* | XY | Assumed | NA | NA |
| Phyllostomidae | *Chiroderma villosum* | XAYAfusion | Known | Blackmon et al. (2019) | NA |
| Phyllostomidae | *Chiroderma vizottoi* | XY | Assumed | NA | NA |
| Phyllostomidae | *Choeroniscus godmani* | XY | Known | Blackmon et al. (2019) | NA |
| Phyllostomidae | *Choeroniscus minor* | XY | Known | Baker et al. (1982) | NA |
| Phyllostomidae | *Choeroniscus periosus* | XY | Assumed | NA | NA |
| Phyllostomidae | *Choeronycteris mexicana* | XY | Assumed | NA | NA |
| Phyllostomidae | *Chrotopterus auritus* | XY | Known | Baker et al. (1982) | NA |
| Phyllostomidae | *Dermanura anderseni* | XY | Assumed | NA | Considered a variant sex chromosome species in “Parsimonious Karyotype” dataset. |
| Phyllostomidae | *Dermanura azteca* | XAfusion | Known | Webster and Jones (1982) | NA |

| | | | | | |
|---|---|---|---|---|---|
| Phyllostomidae | *Dermanura bogotensis* | XY | Assumed | NA | Considered a variant sex chromosome species in "Parsimonious Karyotype" dataset. |
| Phyllostomidae | *Dermanura cinerea* | XAYAfusion | Known | Santos and Souza (1998) | NA |
| Phyllostomidae | *Dermanura glauca* | XY | Assumed | NA | Considered a variant sex chromosome species in "Parsimonious Karyotype" dataset. |
| Phyllostomidae | *Dermanura gnoma* | XY | Assumed | NA | Considered a variant sex chromosome species in "Parsimonious Karyotype" dataset. |
| Phyllostomidae | *Dermanura incomitatus* | XY | Assumed | NA | Considered a variant sex chromosome species in "Parsimonious Karyotype" dataset. |
| Phyllostomidae | *Dermanura phaeotis* | XAYAfusion | Known | Hsu et al. (1968) | NA |
| Phyllostomidae | *Dermanura rava* | XY | Assumed | NA | Considered a variant sex chromosome species in "Parsimonious Karyotype" dataset. |
| Phyllostomidae | *Dermanura rosenbergi* | XY | Assumed | NA | Considered a variant sex chromosome species in "Parsimonious Karyotype" dataset. |
| Phyllostomidae | *Dermanura tolteca* | XAfusion | Known | Hsu et al. (1968) | NA |
| Phyllostomidae | *Dermanura watsoni* | XAYAfusion | Known | Blackmon et al. (2019) | NA |

| | | | | | |
|---|---|---|---|---|---|
| Phyllostomidae | *Desmodus draculae* | XY | Assumed | NA | NA |
| Phyllostomidae | *Desmodus rotundus* | XY | Assumed | NA | NA |
| Phyllostomidae | *Diaemus youngii* | XY | Known | Blackmon et al. (2019) | NA |
| Phyllostomidae | *Diphylla ecaudata* | XY | Assumed | NA | NA |
| Phyllostomidae | *Dryadonycteris capixaba* | XY | Assumed | NA | NA |
| Phyllostomidae | *Ectophylla alba* | XY | Known | Blackmon et al. (2019) | NA |
| Phyllostomidae | *Enchisthenes hartii* | XY | Assumed | NA | NA |
| Phyllostomidae | *Erophylla bombifrons* | XY | Known | Blackmon et al. (2019) | NA |
| Phyllostomidae | *Erophylla sezekorni* | XY | Known | Blackmon et al. (2019) | NA |
| Phyllostomidae | *Gardnerycteris crenulata* | XY | Known | Baker et al. (1982) | NA |
| Phyllostomidae | *Gardnerycteris koepckeae* | XY | Assumed | NA | NA |
| Phyllostomidae | *Glossophaga commissarisi* | XY | Assumed | NA | NA |
| Phyllostomidae | *Glossophaga leachii* | XY | Assumed | NA | NA |
| Phyllostomidae | *Glossophaga longirostris* | XY | Assumed | NA | NA |
| Phyllostomidae | *Glossophaga morenoi* | XY | Assumed | NA | NA |
| Phyllostomidae | *Glossophaga soricina* | XY | Known | Blackmon et al. (2019) | NA |
| Phyllostomidae | *Glyphonycteris behnii* | XY | Assumed | NA | NA |
| Phyllostomidae | *Glyphonycteris daviesi* | XY | Known | Baker et al. (1982) | NA |
| Phyllostomidae | *Glyphonycteris sylvestris* | XY | Assumed | NA | NA |
| Phyllostomidae | *Hsunycteris cadenai* | XY | Assumed | NA | NA |
| Phyllostomidae | *Hsunycteris pattoni* | XY | Assumed | NA | NA |
| Phyllostomidae | *Hsunycteris thomasi* | XY | Assumed | NA | NA |

| | | | | | |
|---|---|---|---|---|---|
| Phyllostomidae | *Hylonycteris underwoodi* | XY | Assumed | NA | NA |
| Phyllostomidae | *Lampronycteris brachyotis* | XY | Known | Baker et al. (1982) | NA |
| Phyllostomidae | *Leptonycteris curasoae* | XY | Assumed | NA | NA |
| Phyllostomidae | *Leptonycteris nivalis* | XY | Assumed | NA | NA |
| Phyllostomidae | *Leptonycteris yerbabuenae* | XY | Assumed | NA | NA |
| Phyllostomidae | *Lichonycteris obscura* | XY | Assumed | NA | NA |
| Phyllostomidae | *Lionycteris spurrelli* | XY | Known | Baker et al. (1982) | NA |
| Phyllostomidae | *Lonchophylla bokermanni* | XY | Assumed | NA | NA |
| Phyllostomidae | *Lonchophylla chocoana* | XY | Assumed | NA | NA |
| Phyllostomidae | *Lonchophylla concava* | XY | Assumed | NA | NA |
| Phyllostomidae | *Lonchophylla dekeyseri* | XY | Assumed | NA | NA |
| Phyllostomidae | *Lonchophylla fornicata* | XY | Assumed | NA | NA |
| Phyllostomidae | *Lonchophylla handleyi* | XY | Assumed | NA | NA |
| Phyllostomidae | *Lonchophylla hesperia* | XY | Assumed | NA | NA |
| Phyllostomidae | *Lonchophylla inexpectata* | XY | Assumed | NA | NA |
| Phyllostomidae | *Lonchophylla mordax* | XY | Assumed | NA | NA |
| Phyllostomidae | *Lonchophylla orcesi* | XY | Assumed | NA | NA |
| Phyllostomidae | *Lonchophylla orienticollina* | XY | Assumed | NA | NA |
| Phyllostomidae | *Lonchophylla peracchii* | XY | Assumed | NA | NA |
| Phyllostomidae | *Lonchophylla robusta* | XY | Known | Baker et al. (1982) | NA |
| Phyllostomidae | *Lonchophylla thomasi* | XY | Known | Blackmon et al. (2019) | NA |
| Phyllostomidae | *Lonchorhina aurita* | XY | Known | Baker et al. (1982) | NA |

| | | | | | |
|---|---|---|---|---|---|
| Phyllostomidae | *Lonchorhina fernandezi* | XY | Assumed | NA | NA |
| Phyllostomidae | *Lonchorhina inusitata* | XY | Assumed | NA | NA |
| Phyllostomidae | *Lonchorhina marinkellei* | XY | Assumed | NA | NA |
| Phyllostomidae | *Lonchorhina orinocensis* | XY | Assumed | NA | NA |
| Phyllostomidae | *Lophostoma brasiliense* | XY | Known | Baker et al. (1982) | NA |
| Phyllostomidae | *Lophostoma carrikeri* | XY | Assumed | NA | NA |
| Phyllostomidae | *Lophostoma evotis* | XY | Assumed | NA | NA |
| Phyllostomidae | *Lophostoma kalkoae* | XY | Assumed | NA | NA |
| Phyllostomidae | *Lophostoma occidentale* | XY | Assumed | NA | NA |
| Phyllostomidae | *Lophostoma schulzi* | XY | Known | Baker et al. (1982) | NA |
| Phyllostomidae | *Lophostoma silvicola* | XY | Known | Baker et al. (1982) | NA |
| Phyllostomidae | *Lophostoma yasuni* | XY | Assumed | NA | NA |
| Phyllostomidae | *Macrophyllum macrophyllum* | XY | Assumed | NA | NA |
| Phyllostomidae | *Macrotus californicus* | XY | Known | Blackmon et al. (2019) | NA |
| Phyllostomidae | *Macrotus waterhousii* | XY | Known | Blackmon et al. (2019) | NA |
| Phyllostomidae | *Mesophylla macconnelli* | XAYAfusion | Known | Baker and Hsu (1970) | NA |
| Phyllostomidae | *Micronycteris brosseti* | XY | Assumed | NA | NA |
| Phyllostomidae | *Micronycteris buriri* | XY | Assumed | NA | NA |
| Phyllostomidae | *Micronycteris giovanniae* | XY | Assumed | NA | NA |
| Phyllostomidae | *Micronycteris hirsuta* | XY | Known | Baker et al. (1982) | NA |
| Phyllostomidae | *Micronycteris matses* | XY | Assumed | NA | NA |
| Phyllostomidae | *Micronycteris megalotis* | XY | Known | Blackmon et al. (2019) | NA |

| | | | | | |
|---|---|---|---|---|---|
| Phyllostomidae | *Micronycteris microtis* | XY | Assumed | NA | NA |
| Phyllostomidae | *Micronycteris minuta* | XY | Known | Blackmon et al. (2019) | NA |
| Phyllostomidae | *Micronycteris sanborni* | XY | Assumed | NA | NA |
| Phyllostomidae | *Micronycteris schmidtorum* | XY | Known | Baker et al. (1982) | NA |
| Phyllostomidae | *Micronycteris yatesi* | XY | Assumed | NA | NA |
| Phyllostomidae | *Mimon bennettii* | XY | Known | Baker et al. (1982) | NA |
| Phyllostomidae | *Mimon cozumelae* | XY | Assumed | NA | NA |
| Phyllostomidae | *Monophyllus plethodon* | XY | Assumed | NA | NA |
| Phyllostomidae | *Monophyllus redmani* | XY | Assumed | NA | NA |
| Phyllostomidae | *Musonycteris harrisoni* | XY | Assumed | NA | NA |
| Phyllostomidae | *Neonycteris pusilla* | XY | Assumed | NA | NA |
| Phyllostomidae | *Phylloderma stenops* | XY | Known | Baker et al. (1982) | NA |
| Phyllostomidae | *Phyllonycteris aphylla* | XY | Known | Blackmon et al. (2019) | NA |
| Phyllostomidae | *Phyllonycteris poeyi* | XY | Assumed | NA | NA |
| Phyllostomidae | *Phyllops falcatus* | XAfusion | Known | Greenbaum et al. (1975) | NA |
| Phyllostomidae | *Phyllostomus discolor* | XY | Known | Blackmon et al. (2019) | NA |
| Phyllostomidae | *Phyllostomus elongatus* | XY | Known | Blackmon et al. (2019) | NA |
| Phyllostomidae | *Phyllostomus hastatus* | XY | Known | Blackmon et al. (2019) | NA |
| Phyllostomidae | *Phyllostomus latifolius* | XY | Known | Baker et al. (1982) | NA |
| Phyllostomidae | *Platalina genovensium* | XY | Assumed | NA | NA |
| Phyllostomidae | *Platyrrhinus albericoi* | XY | Assumed | NA | Considered a variant sex chromosome species in “Parsimonious |

| | | | | | |
|---|---|---|---|---|---|
| | | | | | Karyotype" dataset. |
| Phyllostomidae | *Platyrrhinus angustirostris* | XY | Assumed | NA | Considered a variant sex chromosome species in "Parsimonious Karyotype" dataset. |
| Phyllostomidae | *Platyrrhinus aurarius* | XY | Assumed | NA | Considered a variant sex chromosome species in "Parsimonious Karyotype" dataset. |
| Phyllostomidae | *Platyrrhinus brachycephalus* | XY | Assumed | NA | Considered a variant sex chromosome species in "Parsimonious Karyotype" dataset. |
| Phyllostomidae | *Platyrrhinus chocoensis* | XY | Assumed | NA | Considered a variant sex chromosome species in "Parsimonious Karyotype" dataset. |
| Phyllostomidae | *Platyrrhinus dorsalis* | XY | Assumed | NA | NAConsidered a variant sex chromosome species in "Parsimonious Karyotype" dataset. |
| Phyllostomidae | *Platyrrhinus fusciventris* | XY | Assumed | NA | Considered a variant sex chromosome species in "Parsimonious Karyotype" dataset. |
| Phyllostomidae | *Platyrrhinus guianensis* | XY | Assumed | NA | Considered a variant sex chromosome species in "Parsimonious Karyotype" dataset. |

| | | | | | |
|---|---|---|---|---|---|
| Phyllostomidae | *Platyrrhinus helleri* | XY | Assumed | NA | Considered a variant sex chromosome species in "Parsimonious Karyotype" dataset. |
| Phyllostomidae | *Platyrrhinus infuscus* | XY | Assumed | NA | Considered a variant sex chromosome species in "Parsimonious Karyotype" dataset. |
| Phyllostomidae | *Platyrrhinus ismaeli* | XY | Assumed | NA | Considered a variant sex chromosome species in "Parsimonious Karyotype" dataset. |
| Phyllostomidae | *Platyrrhinus lineatus* | XAYAfusion | Known | Baker and Bickham (1980) | While this karyotype has been studied, certain chromosome morphology attributes relevant to our study were not recorded. |
| Phyllostomidae | *Platyrrhinus masu* | XY | Assumed | NA | Considered a variant sex chromosome species in "Parsimonious Karyotype" dataset. |
| Phyllostomidae | *Platyrrhinus matapalensis* | XY | Assumed | NA | Considered a variant sex chromosome species in "Parsimonious Karyotype" dataset. |
| Phyllostomidae | *Platyrrhinus nigellus* | XY | Assumed | NA | Considered a variant sex chromosome species in "Parsimonious Karyotype" dataset. |

| | | | | | |
|---|---|---|---|---|---|
| Phyllostomidae | *Platyrrhinus nitelinea* | XY | Assumed | NA | Considered a variant sex chromosome species in “Parsimonious Karyotype” dataset. |
| Phyllostomidae | *Platyrrhinus recifinus* | XY | Assumed | NA | Considered a variant sex chromosome species in “Parsimonious Karyotype” dataset. |
| Phyllostomidae | *Platyrrhinus umbratus* | XY | Assumed | NA | Considered a variant sex chromosome species in “Parsimonious Karyotype” dataset. |
| Phyllostomidae | *Platyrrhinus vittatus* | XAYAfusion | Known | Varella-Garcia et al. (1989) | NA |
| Phyllostomidae | *Pygoderma bilabiatum* | XY | Assumed | NA | Considered a variant sex chromosome species in “Parsimonious Karyotype” dataset. |
| Phyllostomidae | *Rhinophylla alethina* | XY | Assumed | NA | NA |
| Phyllostomidae | *Rhinophylla fischerae* | XY | Assumed | NA | NA |
| Phyllostomidae | *Rhinophylla pumilio* | XY | Assumed | NA | NA |
| Phyllostomidae | *Scleronycteris ega* | XY | Assumed | NA | NA |
| Phyllostomidae | *Sphaeronycteris toxophyllum* | XY | Assumed | NA | NA |
| Phyllostomidae | *Stenoderma rufum* | XY | Assumed | NA | Considered a variant sex chromosome species in “Parsimonious Karyotype” dataset. |
| Phyllostomidae | *Sturnira angeli* | XY | Assumed | NA | NA |
| Phyllostomidae | *Sturnira aratathomasi* | XY | Assumed | NA | NA |

| | | | | | |
|---|---|---|---|---|---|
| Phyllostomidae | *Sturnira bakeri* | XY | Assumed | NA | NA |
| Phyllostomidae | *Sturnira bidens* | XY | Assumed | NA | NA |
| Phyllostomidae | *Sturnira bogotensis* | XY | Assumed | NA | NA |
| Phyllostomidae | *Sturnira burtonlimi* | XY | Assumed | NA | NA |
| Phyllostomidae | *Sturnira erythromos* | XY | Assumed | NA | NA |
| Phyllostomidae | *Sturnira hondurensis* | XY | Assumed | NA | NA |
| Phyllostomidae | *Sturnira koopmanhilli* | XY | Assumed | NA | NA |
| Phyllostomidae | *Sturnira lilium* | XY | Assumed | NA | NA |
| Phyllostomidae | *Sturnira ludovici* | XY | Assumed | NA | NA |
| Phyllostomidae | *Sturnira luisi* | XY | Assumed | NA | NA |
| Phyllostomidae | *Sturnira magna* | XY | Assumed | NA | NA |
| Phyllostomidae | *Sturnira mistratensis* | XY | Assumed | NA | NA |
| Phyllostomidae | *Sturnira mordax* | XY | Assumed | NA | NA |
| Phyllostomidae | *Sturnira nana* | XY | Assumed | NA | NA |
| Phyllostomidae | *Sturnira oporaphilum* | XY | Assumed | NA | NA |
| Phyllostomidae | *Sturnira parvidens* | XY | Assumed | NA | NA |
| Phyllostomidae | *Sturnira paulsoni* | XY | Assumed | NA | NA |
| Phyllostomidae | *Sturnira perla* | XY | Assumed | NA | NA |
| Phyllostomidae | *Sturnira sorianoi* | XY | Assumed | NA | NA |
| Phyllostomidae | *Sturnira tildae* | XY | Assumed | NA | NA |
| Phyllostomidae | *Tonatia bidens* | XY | Known | Blackmon et al. (2019) | NA |
| Phyllostomidae | *Tonatia saurophila* | XY | Assumed | NA | NA |
| Phyllostomidae | *Trachops cirrhosus* | XY | Known | Baker et al. (1982) | NA |
| Phyllostomidae | *Trinycteris nicefori* | XY | Known | Baker et al. (1982) | NA |
| Phyllostomidae | *Uroderma bakeri* | XY | Assumed | NA | Considered a variant sex chromosome species in |

| | | | | | |
|---|---|---|---|---|---|
| | | | | | “Parsimonious Karyotype” dataset. |
| Phyllostomidae | *Uroderma bilobatum* | XAYAfusion | Known | Blackmon et al. (2019) | NA |
| Phyllostomidae | *Uroderma magnirostrum* | XAYAfusion | Known | Pieczarka et al. (2013) | NA |
| Phyllostomidae | *Vampyressa elisabethae* | XY | Assumed | NA | Considered a variant sex chromosome species in “Parsimonious Karyotype” dataset. |
| Phyllostomidae | *Vampyressa melissa* | XY | Known | Blackmon et al. (2019) | NA |
| Phyllostomidae | *Vampyressa pusilla* | XAYAfusion | Known | Blackmon et al. (2019) | NA |
| Phyllostomidae | *Vampyressa sinchi* | XY | Assumed | NA | Considered a variant sex chromosome species in “Parsimonious Karyotype” dataset. |
| Phyllostomidae | *Vampyressa thyone* | XAYAfusion | Known | Baker (1973) | NA |
| Phyllostomidae | *Vampyriscus bidens* | XAYAfusion | Known | Blackmon et al. (2019) | NA |
| Phyllostomidae | *Vampyriscus brocki* | XAYAfusion | Known | Blackmon et al. (2019) | NA |
| Phyllostomidae | *Vampyriscus nymphaeus* | XY | Known | Blackmon et al. (2019) | NA |
| Phyllostomidae | *Vampyrodes caraccioli* | XAYAfusion | Known | Baker (1973) | NA |
| Phyllostomidae | *Vampyrum spectrum* | XY | Known | Baker et al. (1982) | NA |
| Phyllostomidae | *Xeronycteris vieirai* | XY | Assumed | NA | NA |

Supplemental Table 2: Phylogenetic logistic regression p-values (upper, corrected for false discovery rate) and estimates (lower) from correlation testing between ecoclimatic variables and the presence of sex-autosome fusions in “known karyotype” species only.

| Family (sp. with known karyotypes only) | | Species range size | Absolute latitude | Annual mean temperature | Annual precipitation | Temperature seasonality | Precipitation seasonality | Elevation |
|---|---|---|---|---|---|---|---|---|
| Herpestidae (16 species) | *P* | 0.909 | 0.995 | 0.909 | 0.909 | 0.909 | 0.909 | 0.909 |
| | Est. | 0.0005 | 0.0002 | 0.005 | 0.0009 | 0.0002 | -0.007 | -0.002 |
| Soricidae (49 species) | *P* | 0.903 | 0.810 | 0.903 | 0.810 | 0.810 | 0.974 | 0.903 |
| | Est. | 0.0002 | 0.072 | -0.0007 | -0.0005 | 0.0002 | -0.0003 | 0.0002 |
| Phyllostomidae (71 species) | *P* | 0.994 | 0.994 | 0.994 | 0.994 | 0.994 | 0.994 | 0.994 |
| | Est. | -0.0003 | -0.015 | -0.009 | -0.0001 | -0.000002 | 0.008 | -0.00002 |

Supplemental Table 3: Mean p-values (upper) and estimates (center), and permutation p-values calculated via the formula $1 - [(\Sigma(number\ of\ tests\ with\ pval \leq 0.05) + 1) \div (1000 + 1)]$ (lower, corrected for false discovery rate) from *sensiPhy* analysis over 1000 credible trees for each family, which tested correlation between ecoclimatic variables and sex-autosome fusions in “known karyotype” species only.

| Family (sp. with known karyotypes only) | | Species range size | Absolute latitude | Annual mean temperature | Annual precipitation | Temperature seasonality | Precipitation seasonality | Elevation |
|---|---|---|---|---|---|---|---|---|
| Herpestidae (16 species) | Mean *p* | 0.875 | 0.998 | 0.782 | 0.271 | 0.433 | 0.764 | 0.188 |
| | Est. | 0.0001 | 0.00006 | 0.005 | 0.0009 | 0.0002 | -0.006 | -0.003 |
| | Permutation p-values | 0.999 | 0.999 | 0.999 | 0.999 | 0.999 | 0.999 | 0.999 |
| Soricidae (49 species) | Mean *p* | 0.611 | 0.233 | 0.788 | 0.276 | 0.885 | 0.861 | 0.698 |
| | Est. | 0.0002 | 0.073 | -0.0007 | -0.0007 | 0.00002 | -0.006 | 0.0001 |
| | Permutation p-values | 0.999 | 0.999 | 0.999 | 0.999 | 0.999 | 0.999 | 0.999 |
| Phyllostomidae (71 species) | Mean *p* | 0.673 | 0.690 | 0.676 | 0.740 | 0.994 | 0.680 | 0.318 |
| | Est. | 0.0002 | -0.012 | -0.008 | -0.0001 | <0.000001 | 0.007 | -0.001 |
| | Permutation p-values | 0.999 | 0.999 | 0.999 | 0.999 | 0.999 | 0.999 | 0.999 |

Supplemental Table 4: Phylogenetic logistic regression p-values (upper, corrected for false discovery rate) and estimates (lower) from correlation testing between ecoclimatic variables and the presence of sex-autosome fusions under the “parsimonious karyotype” assumption.

| Family ("parsimonious karyotype") | | Species range size | Absolute latitude | Annual mean temperature | Annual precipitation | Temperature seasonality | Precipitation seasonality | Elevation |
|---|---|---|---|---|---|---|---|---|
| Herpestidae | *P* | 0.625 | 0.997 | 0.997 | 0.625 | 0.625 | 0.997 | 0.625 |
| | Est. | 0.001 | 0.0002 | 0.001 | 0.0005 | 0.0003 | 0.00002 | -0.0009 |
| Phyllostomidae | *P* | 0.964 | 0.964 | 0.964 | 0.964 | 0.964 | 0.964 | 0.964 |
| | Est. | 0.00009 | -0.0004 | -0.005 | -0.00001 | -0.0001 | -0.0003 | 0.00004 |

Supplemental Table 5: Mean p-values (upper) and estimates (center), and permutation p-values calculated via the formula $1 - [(\Sigma(number\ of\ tests\ with\ pval \leq 0.05) + 1) \div (1000 + 1)]$ (lower, corrected for false discovery rate) from *sensiPhy* analysis over 1000 credible trees, which tested correlation between ecoclimatic variables and sex-autosome fusions under the "parsimonious karyotype" assumption.

| Family ("parsimonious karyotype") | | Species range size | Absolute latitude | Annual mean temperature | Annual precipitation | Temperature seasonality | Precipitation seasonality | Elevation |
|---|---|---|---|---|---|---|---|---|
| Herpestidae | Mean *p* | 0.359 | 0.895 | 0.908 | 0.365 | 0.262 | 0.864 | 0.631 |
| | Est. | 0.001 | -0.003 | 0.001 | 0.0005 | 0.0003 | -0.001 | -0.0004 |
| | Permutation p-values | 0.999 | 0.999 | 0.999 | 0.999 | 0.999 | 0.999 | 0.999 |
| Phyllostomidae | Mean *p* | 0.663 | 0.939 | 0.379 | 0.713 | 0.333 | 0.903 | 0.789 |
| | Est. | 0.00009 | -0.0002 | -0.005 | -0.00005 | -0.0001 | -0.0002 | 0.00003 |
| | Permutation p-values | 0.999 | 0.999 | 0.999 | 0.999 | 0.999 | 0.999 | 0.999 |
